# Epigenetic control of commensal induced Th2 Responses and Intestinal immunopathology

**DOI:** 10.1101/2024.08.30.610485

**Authors:** Kishan A. Sangani, Morgan E. Parker, Hope D. Anderson, Li Chen, Surya P. Pandey, Joseph F. Pierre, Marlies Meisel, Samantha J. Riesenfeld, Reinhard Hinterleitner, Bana Jabri

## Abstract

Understanding the initiation of T-helper (Th)-2 immunity is crucial for addressing allergic diseases that have been linked to the commensal microbiota. However, Th2 responses are notably absent from known host-microbiota intestinal immune circuits. Notably, the commensal protist *Tritrichomonas* induces a transient innate ILC2 circuit rather than a chronic Th2 circuit. Canonical Th2 responses rely on the induction of IL-4 production by innate cells. This study shows that the absence of *Tet2*, a DNA demethylase, reprograms naïve T cells to autonomously produce IL-4 upon T cell receptor stimulation, bypassing the need for IL-4 from innate cells for Th2 differentiation. Loss of this checkpoint induces chronic Th2 responses to *Tritrichomonas*, associated with IL-25-dependent barrier dysfunction and increased susceptibility to allergic pathology in response to dietary antigens.

**Sentence Summary:** Regulation of cell autonomous IL-4 in T cells is critical to prevent dysregulated Th2 immunity to commensals and predisposition to allergy.

## Main Text

The prevalence of type 2 immune-mediated disorders, such as food allergies, has seen a significant increase in developing countries over the past few decades^1,2^. While it is well recognized that both genetic and environmental factors contribute to an individual’s predisposition to type-2 immune dysregulation^3,4^, the specific mechanisms underlying this predisposition remain elusive. Classically, the differentiation of T-helper (Th) responses necessitates antigen presentation in the presence of polarizing cytokines. However, Th2 responses stand apart in that their polarizing cytokine, interleukin (IL)-4, is not typically produced by professional antigen-presenting cells^5,6^. Paradoxically, IL-4 also serves as the primary effector cytokine for the Th2 lineage^7^, which has posed challenges in identifying the initial source of IL-4 in models of type-2 inflammation. An indirect or decentralized model for type-2 immunity has been proposed, which involves the orchestration of antigen presentation by dendritic cells in the presence of IL-4 derived from granulocytes or innate-like lymphocytes^5,8^. Notably, unlike for helminths or other type 2 immune challenges^9,10^, there are no reports of commensal microbes inducing significant IL-4 production by innate immune cells. This may explain the absence of homeostatic Th2 responses at mucosal barrier sites, such as the intestine, where Th1 and Th17 responses to the microbiota are prevalent^11–13^. Commensal protists of the Tritrichomonas species have been instead shown to trigger a type-2 innate immune circuit in the small intestine that involves the activation of tuft cells and the production of IL-25, which, in turn, stimulates type 2 innate lymphoid cells (ILC2s) to produce IL-5 and IL-13^14–17^. This type 2 circuit resembles what is known to occur in the context of helminth infections, but strikingly lacks the participation of adaptive Th2 cells^2,17–20^, which are essential for helminth clearance^2,18^. Interestingly, this activation is transient despite the persistence of Tritrichomonas^14^, suggesting that we have evolved mechanisms to prevent the establishment of chronic type-2 immunity in the gut, which could be detrimental to health.

DNA methylation at CpG motifs is an important epigenetic modification implicated in gene regulation and expression^21^. The characterization of specific DNA methylase and demethylase enzymes has enabled study of specific mechanisms and contexts in which DNA methylation is an important determinant of immune cell fate and differentiation^22^. These enzymes are often mutated in hematologic malignancies and the demethylase TET2 is among the most frequently mutated, underscoring important functions in hematopoietic differentiation and immune function^23–25^. In a previous study, we reported that microbiota-dependent intestinal barrier dysfunction was responsible for the development of preleukemic myeloproliferation in aged *Tet2*-deficient mice, with the jejunum being the primary intestinal site affected by the absence of *Tet2* expression as assessed by transcriptional changes^26^. Furthermore, while we identified commensal Gram-positive bacteria as sufficient to promote preleukemic myeloproliferation^26^, the specific microbial origin responsible for barrier dysfunction in *Tet2*-deficient mice has remained elusive. In this study, we aimed to uncover the mechanisms underlying small intestinal barrier dysfunction in *Tet2*-deficient mice and to explore the potential associated immunopathological implications.

## Chronic jejunal IL-25 signaling in the absence of *Tet2*

We first examined our intestinal gene expression dataset for enrichment of published cell identity signatures^27^. We found that among nearly 130 cell signatures, the only one significantly enriched in the jejunum (false discovery rate (FDR) < 0.05) was that of tuft cells (Fig. 1A), while other intestinal segments did not exhibit significant enrichment for any cell signatures (Fig. S1A).

**Figure 1:**
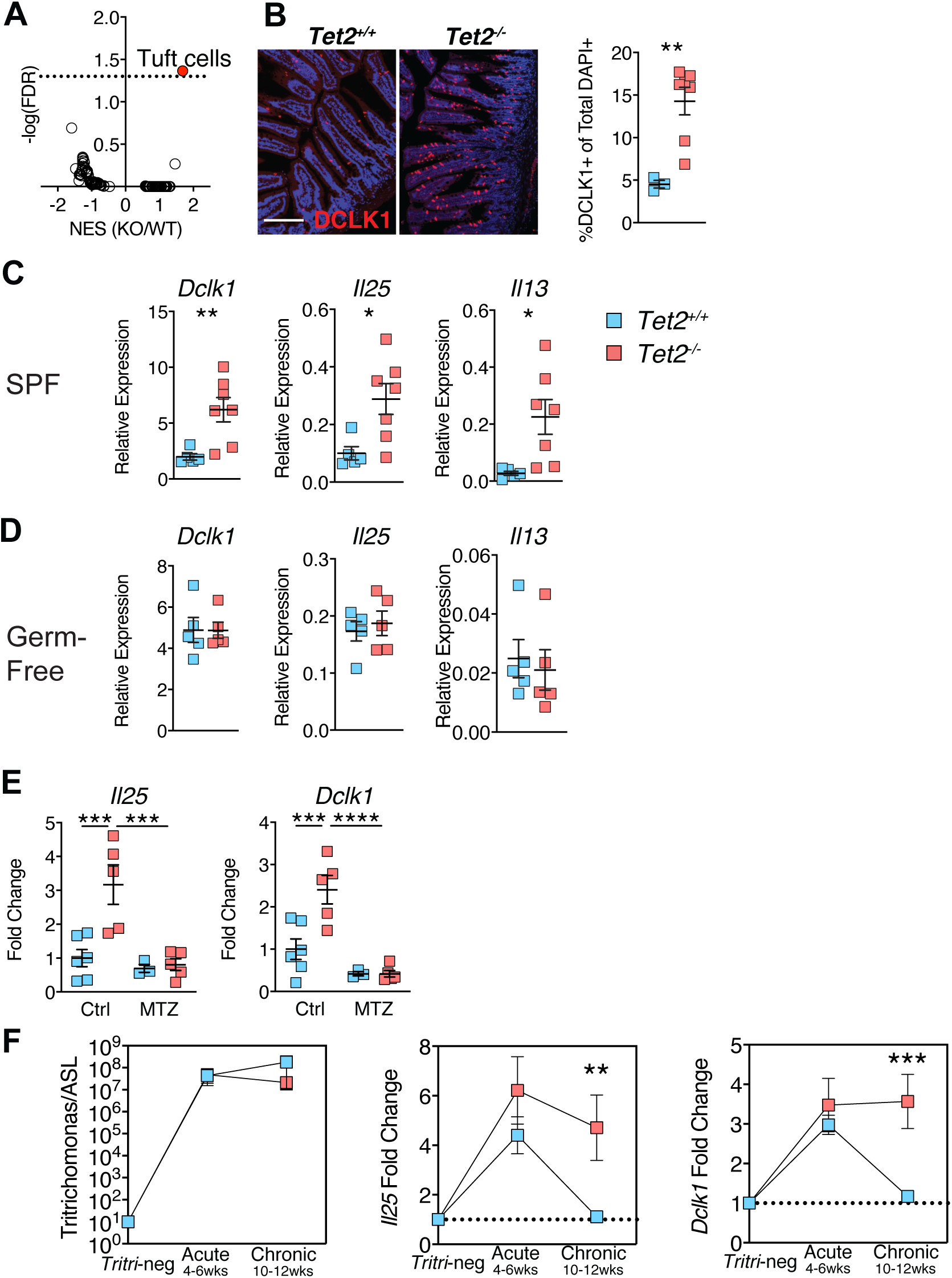
Chronic Tritrichomonas-dependent Tuft-IL25 signature in the absence of Tet2. A, Volcano plot of normalized enrichment scores and adjusted p-values for various cell signatures from normalized RNAseq expression data from Tet2+/+ and Tet2-/- jejunum tissue. B, OCT fixed segments of jejunum were sectioned and stained with DLCK1 and mounted with DAPI and proportions of DAPI+ and DLCK1+ cells were quantified in the epithelium. C/D, Quantitative PCR of jejunal segments from SPF and Germ-free Tet2+/+ and Tet2-/- mice expressed relative to *Gapdh*. E, Quantitative PCR of Tritrichomonas relative to IL-25 circuit genes relative to *Gapdh* in the jejunum from Tet2+/+ and Tet2-/- mice treated with metronidazole (MTZ) or control water for 4 weeks. IL-25 circuit genes expressed as fold-change relative to expression in control treated wildtype jejunum. F, Quantitative PCR of Tritrichomonas relative to 16s in cecal contents (left) or *Il25* and *Dclk1* relative to *Gapdh* in jejunal tissue (middle, right) from Tet2+/+ and Tet2-/- mice colonized with Tritrichomonas acutely (4-6 weeks) or chronically (10-12 weeks). *Il25* and *Dclk1* represented as fold change over relative expression in jejunum from uncolonized Tet2+/+ mice. Dotted line indicates baseline fold change of 1. Statistical analysis was performed by two-sided Student’s t-test in (B-D) one-way ANOVA in (E,F) with Tukey multiple-comparison test. p-values: * (<0.05) **(<0.01) *** (<0.001). All panels representative of at least 2 independent experiments.

We confirmed the expansion of tuft cells through immunofluorescent staining for the tuft cell marker Dclk1 (Fig. 1B) and quantitative polymerase chain reaction (qPCR) (Fig. 1C). Additionally, in line with previous reports^19^, the expansion of tuft cells was associated with an upregulation of *Il25* and *Il13* expression (Fig. 1C) and a lengthening of the small intestine (Fig. S1B). We also observed expansion of the goblet cell lineage as evidenced by periodic acid shift (PAS) staining and RNA expression of the goblet cell marker *Spink4* (Fig. S1C-D).

While we had previously established that the microbial signal for barrier dysfunction was distinct from the one required for preleukemic myeloproliferation, we were unable to identify the specific microbe(s) responsible for the development of barrier dysfunction in *Tet2*-deficient mice^26^. We first asked if this circuit was microbial-dependent using germ-free mice and indeed found the tuft cell – type II immunity circuit hallmark genes were no longer differentially expressed in these conditions (Fig. 1D, S1C). We next confirmed that the tuft-IL-25-IL-13 signature observed in specific pathogen-free (SPF) *Tet2*^-/-^ mice was dependent on the presence of microbes by treating SPF *Tet2*^-/-^ mice with broad-spectrum antibiotics. As expected, approximately 99% (1376/1395) of the DEGs (p-adj < 0.05) were lost upon antibiotic treatment (Fig. S1E). Notably, the enrichment of the tuft cell signature in *Tet2*^-/-^ mice was also lost (Fig. S1F).

Drawing from reports that associate *Tritrichomonas* with the tuft-IL-25-IL-13 circuit^14,15,20,28^, we hypothesized that *Tritrichomonas* was responsible for driving the dysregulated activation of this circuit in *Tet2*^-/-^ mice. We confirmed the presence of these species in our colony by conducting ribosomal 28S internal transcriber sequencing (ITS) on cecal contents and phylogenetic comparison to published sequences of *Tritrichomonas* 28S (Fig. S1G). Importantly, the identified *Tritrichomonas* species did not show a difference in abundance in *Tet2*^-/-^ mice compared to littermate controls (Fig. S1H). Finally, treatment with metronidazole, which clears *Tritrichomonas*^29^, reversed the upregulation of *Il25* and *Dclk1* in *Tet2*^-/-^ mice (Fig. 1E). Together, these data suggested the absence of *Tet2* altered the host response to *Tritrichomonas* rather than its colonization and led to the establishment of a persistent tuft-IL-25 circuit.

To demonstrate that *Tritrichomonas* was the causative factor behind this observed phenotype, we conducted colonization studies using purified protists. In line with previous observations showing that the type 2 immune signaling circuit induced by succinate-producing protists is transient^14–16^, wild-type mice colonized with *Tritrichomonas* at weaning exhibited an initial acute induction followed by a subsequent chronic regression of *Il25* expression (Fig. 1F, S1I). This pattern was also evident in the changes in *Dclk1* and *Spink4* expression, which are typically found in tuft cells and goblet cells (Fig. 1F, S1I). Notably, the decrease in *Il25* occurred despite the persistent presence of *Tritrichomonas* within the commensal microbiota (as illustrated in Fig. 1F). In contrast, *Tet2^-/-^*mice with similar levels of *Tritrichomonas* (Fig. 1F) did not downregulate *Il25*, *Dclk1* (Fig. 1F) or *Spink4* (Fig. S1I), even up to 10-12 weeks post-colonization. These findings collectively demonstrate that the protist *Tritrichomonas* is necessary and sufficient to initiate a chronic type-2 immune program in the small intestine of *Tet2^-/-^* mice, while wild-type mice only induce a transient program. Importantly, the difference in the host response observed in *Tet2^-/-^* mice cannot be attributed to differences in *Tritrichomonas* colonization or persistence.

## *Tet2*-deficiency redirects an ILC2 driven type 2 circuit to Th2 cells

Previous studies have shown that in wild-type mice, the increase in tuft-cell differentiation upon protist colonization is present in *Rag2^-/-^*but not *Rag2^-/-^ Il2rγ^-/-^* mice and that ILC2 activation is associated with *Tritrichomonas* colonization^14^. Accordingly, colonization with *Tritrichomonas* resulted in increased Ki67 expression in ILC2s from mice of both genotypes (Fig. 2A). The sustained nature of the type-2 immune response, in conjunction with the continued presence of *Tritrichomonas*, prompted our hypothesis that the absence of *Tet2* may be associated with establishment of an adaptive Th2 response to the protist rather than a persistent innate response. Consistent with this hypothesis, *Tritrichomona*s colonization in *Tet2^-/-^*mice led to a significant increase in the proportion of Gata3^+^ Foxp3^-^ CD4^+^ among CD4 T cells in the *lamina propria* (Fig. 2B-C). Treatment with metronidazole, which eliminates the protist^29^ (Fig. S2C), resulted in reduced activation of Th2 cells (Fig, 2D), as assessed by CD69 expression^30^, suggesting that the Th2 subset was induced in response to *Tritrichomona*s.

**Figure 2:**
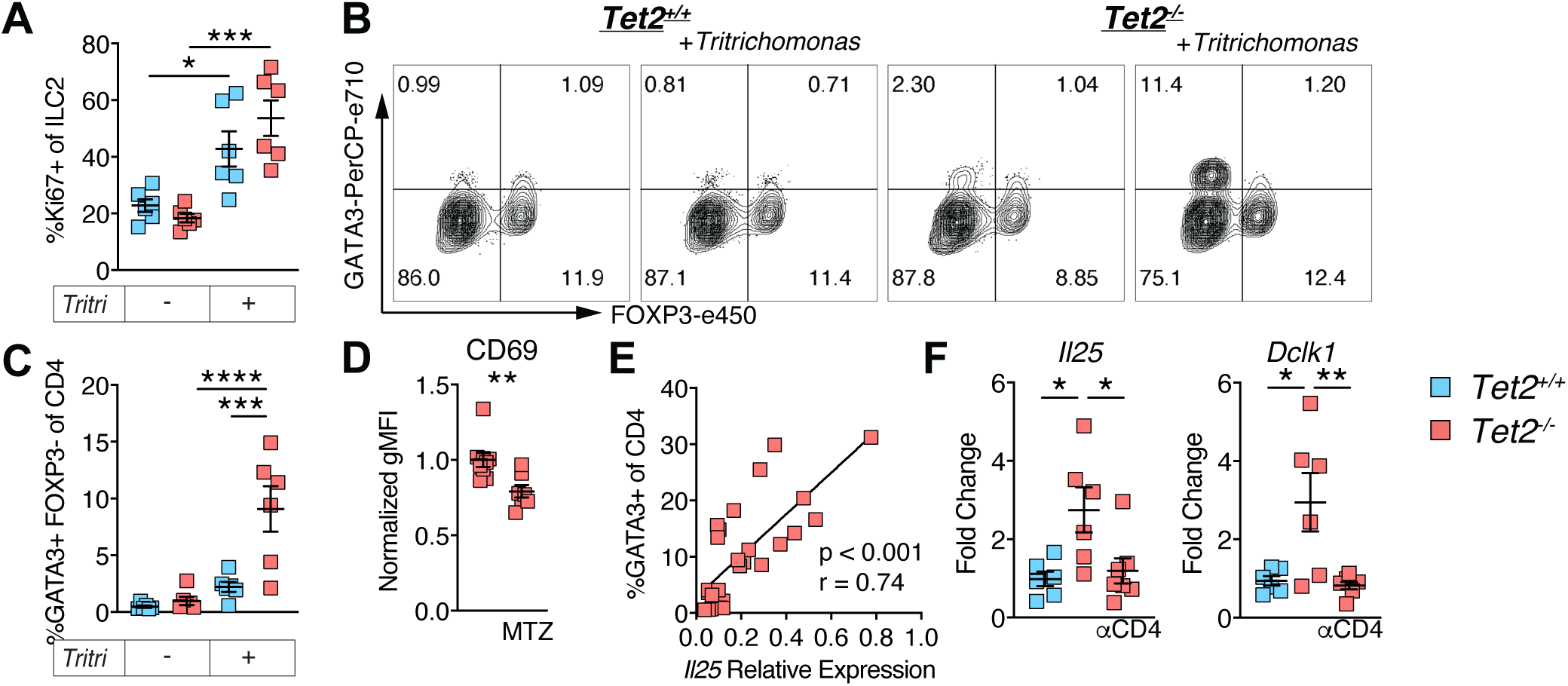
Tet2-deficient Th2 cells are induced after Tritrichomonas colonization and propagate the chronic IL25 signaling circuit. A, Proportion of Ki67+ ILC2s (Live CD45+ Lin-CD90+ KLRG1+ GATA3+) in the jejunum lamina propria of uncolonized mice or after 4 weeks of *Tritrichomonas*-colonization in Tet2+/+ and Tet2-/- mice. B, Representative FOXP3 and GATA3 staining of jejunum lamina propria CD45+ Epcam-Ter119-Live CD90+ CD4+ CD8a-cells from uncolonized mice or after 4 weeks of *Tritrichomonas*-colonization in Tet2+/+ and Tet2-/- mice. C, Proportion of FOXP3-GATA3+ of CD4 T cells from the jejunal lamina propria of uncolonized and 4-week *Tritrichomonas*-colonized Tet2+/+ and Tet2-/- mice. D, Geometric mean fluorescent intensity of CD69 on Foxp3-GATA3+ CD4 T cells from the jejunal lamina propria of *Tritrichomonas*-colonized Tet2-/- mice treated with 1g/L metronidazole or control water for 4 weeks. E, Paired relative expression of *Il25* in the jejunum of Tet2-/- mice along with the proportion of Foxp3-Gata3+ of CD4 T cells in the jejunum lamina propria. F, Quantitative PCR of IL-25 circuit genes relative to Gapdh in the jejunum of Tet2+/+ and Tet2-/- mice chronically colonized with *Tritrichomonas* and treated with isotype or CD4-depleting antibody. Represented as fold change over Tet2+/+ mice. Statistical analysis was performed by one-way ANOVA in (A,C,F) with Tukey multiple-comparison test, two-sided Student’s t-test in (D), and Pearson correlation in (E). p-values: * (<0.05) **(<0.01) *** (<0.001) **** (<0.0001). All panels are representative of 2-3 independent experiments.

In contrast, and as previously shown^14,16,17^, littermate wild-type control mice failed to develop a predominant Th2 response (Fig. 2B-C). Interestingly, while colonization with *Tritrichomona*s was associated with a relative decrease in Th17 and Foxp3^+^ regulatory T cells (Fig S2A-B), this decrease was not influenced by *Tet2.* No changes were observed in the proportion of Th1 cells among CD4 cells or ILC2s within the *lamina propria* (Fig S2A-B). Taken together, these data suggest the absence of *Tet2* allows for the establishment of an adaptive Th2 response to a commensal protist, whereas in wildtype mice, this response remains primarily restricted to the ILC2 compartment.

We next assessed the relationship between this adaptive Th2 population and chronic IL25-circuit activation in *Tet2*-deficient mice. Jejunal *Il25* expression in *Tet2^-/-^* mice chronically colonized with *Tritrichomonas* was significantly correlated with the frequency of Th2 cells in the *lamina propria* (Fig 2E). To ascertain the necessity of Th2 cells in the persistent upregulation of IL-25, we initially confirmed the requirement of the adaptive immune system by evaluating the expression of *Il25* in *Tet2*-sufficient and -deficient mice crossed onto a *Rag2*-deficient background (Fig S2D). Next, we conducted a more targeted two-week depletion of CD4+ lymphocytes in mice chronically colonized with *Tritrichomonas* using a neutralizing antibody. The results revealed a notable decrease in both *Il25* and *Dclk1* expression, establishing that Th2 lymphocytes were critical in propagating the chronic IL-25 signaling circuit in *Tet2*-deficient mice in the presence of *Tritrichomonas* (Fig 2F).

## CD4 T-cell autonomous IL-4 production drives Th2 differentiation in the absence of *Tet2*

We next wanted to identify the specific cell type in which *Tet2* was necessary to prevent the initiation of a Th2 response following *Tritrichomonas* colonization. To this end, we generated crosses between *Tet2*-floxed mice and various Cre-driver mouse lines, including those targeting hematopoietic cells (*Vav*-Cre), lymphocyte lineages (*hCD2*-Cre), intestinal epithelial cells (*villin (Vil)*-Cre), myeloid cells (*LysM*-Cre), and ILC2 cells (*Il5*-Cre). The combined findings pointed to a connection between the dysregulated IL-25-Th2 circuit and *Tet2* deficiency within the lymphocyte compartment (*hCD2*-Cre) (Fig. S3A-C, Fig 3A). More specifically, the deletion of *Tet2* in T cells, using the *Cd4*-cre driver, was found to be sufficient for the development of Th2 cells and chronic upregulation of *Il25* in response to *Tritrichomonas* colonization (Fig 3B-C).

**Figure 3:**
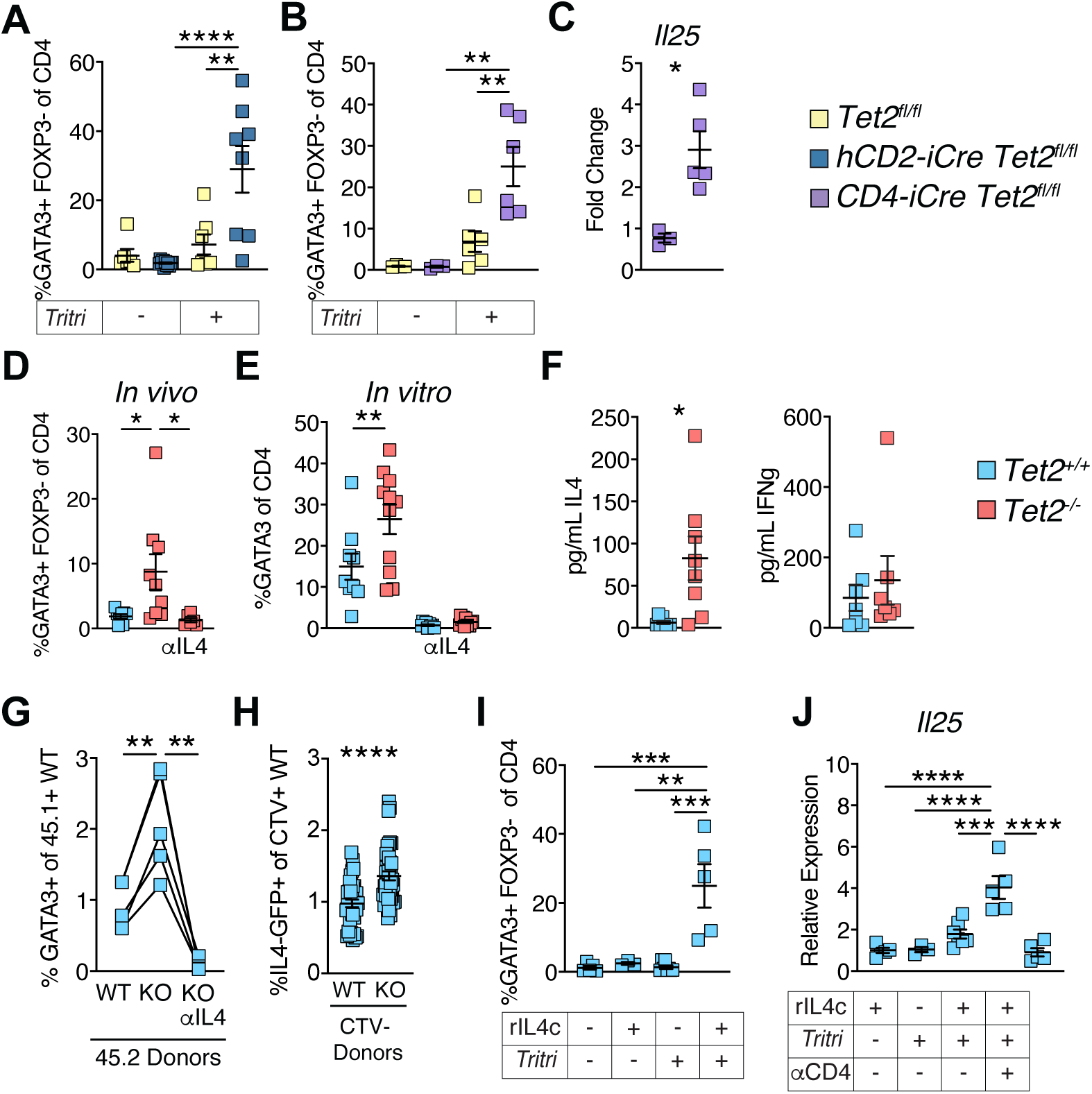
*Tet2* is a CD4-intrinsic checkpoint for IL-4 production and Th2 differentiation. A/B, Proportion of FOXP3-GATA3+ of CD4 T cells from the jejunal lamina propria of uncolonized and 6-week *Tritrichomonas*-colonized mice of indicated genotypes. C, Quantitative PCR of *Il25* relative to *Gapdh* in the jejunum of uncolonized and Tritrichomonas-colonized CD4-Cre+ Tet2 fl/fl mice. Expressed as fold change over uncolonized littermate Tet2 fl/fl mice. D, Proportion of Foxp3-GATA3+ of CD4 T cells from the jejunal lamina propria of Tritrichomonas-colonized Tet2+/+ and Tet2-/- mice treated with anti-IL4 or isotype control antibody for 2 weeks after colonization. E, Proportion of FOXP3-GATA3+ of cultured Naïve CD4 T cells of indicated genotypes cultured for 4 days in non-polarizing conditions (anti-CD3e, anti-CD28 and recombinant IL-2) with and without anti-IL4. F, ELISA quantification of IL-4 (left) and IFNg (right) in the supernatant of naïve CD4 cells cultured for 4 days in (E). G, FOXP3-GATA3+ proportion of cultured CD45.1 Tet2+/+ Naïve CD4 T cells cultured at a 1:4 ratio with Tet2+/+ or Tet2-/- Naïve CD4 T cells for 4 days in non-polarizing conditions with and without anti-IL4. H, Proportion of GFP expressing cells of the CTV-labeled 4Get Naïve CD4 T cells cultured at a 1:4 ratio with Tet2+/+ or Tet2-/- Naïve CD4 T cells in non-polarizing conditions for 4 days. I, Proportion of FOXP3-GATA3+ of CD4 T cells from the jejunal lamina propria of Tritrichomonas-colonized and uncolonized Tet2+/+ injected weekly with rIL4c or vehicle for 4 weeks. J, Quantitative PCR of *Il25* relative to *Gapdh* in the jejunum of Tritrichomonas-colonized and uncolonized Tet2+/+ injected weekly with rIL4c, vehicle, or anti-CD4 as indicated for 4 weeks. Statistical analysis was performed by two-sided Student’s t-test in (C, F), repeated measures one-way ANOVA with Dunnet’s multiple comparison test in (G), paired t-test in (H) and one-way ANOVA with Tukey multiple-comparison test in (A, B, D, E, I, J). p-values: * (<0.05) **(<0.01) *** (<0.001) **** (<0.0001). All panels representative of at least 2 independent experiments.

Having found that *Tet2* deficiency within T cells resulted in a Th2 response to *Tritrichomonas*, we next sought to define the mechanism underlying this aberrant response. IL-4 is a prototypical cytokine for Th2 cell differentiation^7^, and blocking IL-4 during protist colonization, prevented the induction of Th2 cells in the small intestine of *Tet2^-/-^* mice (Fig 3D). This established that development of dysregulated Th2 responses to the commensal protist *Tritrichomonas* in *Tet2^-/-^* mice required IL-4. We then aimed to investigate whether *Tet2* also controlled Th2 differentiation or function in other contexts where IL-4 is implicated. After infection with the helminth *Strongyloides venezuelensis*, *Tet2*-deficient mice and their littermate controls exhibited no significant difference in Th2 induction or infection clearance (Fig S3D-E). Importantly, this infection model involves IL-4 production by innate immune cells^9^. Furthermore, absence of *Tet2* also had no impact on *in vitro* Th2 differentiation under Th2 culture conditions that were supplied with recombinant IL-4 (Fig S3F). Taken together, these observations demonstrate that under conditions where IL-4 is provided to naïve T cells, the absence of *Tet2* does not affect Th2 differentiation.

Based on these findings, we hypothesized that the differences in Th2 induction after *Tritrichomonas* colonization in *Tet2*-deficient mice were not due to differences in IL-4 signaling but rather to naïve *Tet2*-deficient T cells uniquely gaining the ability to produce IL-4 upon T cell receptor (TCR) stimulation. As shown in Figure 3E, *Tet2*-deficient naïve CD4 T cells under non-polarizing conditions (TCR stimulation in the presence of IL-2 but without an external source of IL-4) showed a significantly heightened differentiation into GATA3^+^ CD4^+^ T cells compared to wild-type cells. Importantly, the capability of *Tet2*-deficient naïve T cells to differentiate into GATA3^+^ T cells under non-polarizing conditions was revealed to be IL-4-dependent, indicating the cell-autonomous IL-4 production was important for Th2 induction (Fig. 3E). Finally, although some wild-type cultures occasionally had GATA3-expressing T cells induced under non-polarizing conditions, only *Tet2*-deficient T cell cultures produced detectable levels of IL-4 (Fig. 3F). These findings suggest *Tet2* regulates IL-4 production by naïve CD4 T cells after TCR stimulation.

If the intrinsic ability of naïve *Tet2*-deficient T cells to produce IL-4 is the decisive factor for Th2 differentiation under non-polarizing conditions, co-culturing wild-type cells with *Tet2*-deficient T cells should enable wild-type T cells to differentiate into Th2 cells in the absence of exogenous IL-4. To investigate this premise, we employed a co-culture system involving *Tet2*-deficient naïve CD4 T cells in the presence of *Tet2*-sufficient naïve CD4 T cells at a 4:1 (KO:WT) ratio (Fig. S3G). An IL-4 dependent increase in GATA3 expression was observed in wild-type naïve T cells only when they were co-cultured with *Tet2*-deficient naïve CD4 T cells (Fig 3G). Additionally, utilizing *Tet2*-sufficient naïve CD4 T cells from bi-cistronic IL-4 reporter 4get mice^31^, we were able to demonstrate that co-cultured wild-type naïve T cells exhibited heightened IL-4-FP expression when in the presence of naïve *Tet2*-deficient CD4 T cells under non-polarizing conditions (Fig. S3G & Fig. 3H). These results suggest that *Tet2* regulates the intrinsic capacity of naive T cells to produce IL-4, eliminating the need for exogenous IL-4 for their differentiation into Th2 cells. In turn, if this is the mechanism by which Th2 immunity is induced after *Tritrichomonas* colonization in *Tet2* deficient mice, then providing a source of IL-4 to wildtype animals should be sufficient for establishing commensal-induced Th2 responses.

To test this hypothesis, we administered exogenous IL-4 complex to *Tet2*-sufficient wild-type mice, both colonized or uncolonized with *Tritrichomonas,* and assessed the presence of Th2 effector populations in the *lamina propria*. Notably, the induction of Th2 cells required both the delivery of the IL-4 complex and *Tritrichomonas* colonization (Fig. 3I). Furthermore, the presence of Th2 cells was essential for significant upregulation of *Il25* in the small intestine (Fig. 3J). These findings emphasize stringent control of intrinsic IL-4 production by naïve T cells prevents the initiation of a chronic Th2-IL25 circuit in response to a commensal protist. When this intrinsic control is lost, *Tritrichomonas* can induce Th2 cells (Fig. 3D), which subsequently propagate a chronic IL-25 signaling circuit (Fig 2E-F, Fig 3J).

## Genomic reorganization in the absence of *Tet2* permits cell-autonomous IL-4 production

To better understand the mechanisms allowing *Tet2*-deficient naive T cells to develop Th2 profiles under non-polarizing conditions, we performed bulk RNA-sequencing (RNA-seq) of naive CD4 T cells from *Vav*-Cre *Tet2*fl/fl and *Vav*-Cre-negative littermates, both before and after varying intervals of TCR stimulation (4h, 24h, 48h, 96h) under non-polarizing conditions as described above. First, we confirmed the acquisition of a robust Th2 program in *Tet2*-deficient cells after 96 hours of TCR stimulation by assessing the expression of *Il4* and *Il13* (Fig S4A) as well as a broad enrichment in targets determined to be positively regulated by GATA3 in Th2 cells, derived from CHIP-seq and *Gata3*-deletion studies (Fig S4B)^32^. While the upregulation of this Th2 program was only observed at the 96-hour timepoint, we noticed that approximately 150 genes were differentially expressed in naïve, unstimulated *Tet2*-deficient CD4 T cells compared to their wildtype counterparts (Fig 4A). Genes more highly expressed in *Tet2-*deficient naïve cells included several critical regulators of Th2 differentiation, such as *Irf4* and *Egr3* (Fig 4A). Furthermore, hypergeometric pathway enrichment analysis of hallmark and gene ontology (GO) gene sets indicated that *Tet2*-deficient naïve T cells displayed a pre-activated state.^33,34^ For instance, NFκB and Stat5 signaling Hallmark pathways, as well as GO pathways related to immune activation were overrepresented in naïve *Tet2*-deficient CD4 T cells (Fig S4C). These findings are particularly relevant given that NFκB-signaling was reported to be crucial for GATA3 upregulation during Th2 differentiation and to cooperate with NFAT to upregulate IL-4 during the activation of Th2 cells^35^. Similarly, STAT5 was shown to collaborate with GATA3 to prime the Th2 lineage^36^. As a result, naïve *Tet2*-deficient cells may represent a pre-poised state that already express hallmarks of Th2 differentiation at the transcriptional level, although they are indistinguishable from naïve *Tet2*-sufficient CD4 T cells by conventional naïve surface markers (Fig S4D). We noticed in our *in vitro* differentiation assay that, although *Tet2*-deficient T cells overall exhibited a significantly increased capacity to differentiate into GATA3^+^ T cells under non-polarizing conditions compared to wild-type T cells, there was a wide range of induction across biological replicates (Fig 3E, S4A). These data are reminiscent of biological variation in Th2 induction after *Tritrichomonas* colonization in *Tet2-*deficient mice (Fig 2C,E). Thus, to more precisely identify the genes and pathways that enable *Tet2*-deficient naïve CD4 T cells to adopt a Th2 program, we divided our *Tet2*-deficient samples based on their final expression of *Gata3*, termed Gata3^hi^ or Gata3^lo^ (Fig S4A). We then compared their expression profiles to those of wild-type T cells at multiple time points before and after stimulation. Genes upregulated in naïve *Tet2*-deficient CD4 T cells within the Gata3^hi^ group compared to wildtype, but not the Gata3^lo^ group compared to wildtype, included components of the previously referenced NFκB signaling and STAT5 signaling pathways (*Tnfaip3*, *Nr4a3*, *Tnfrsf4*, *Irf4*), as well as broader regulators of transcription such as the histone modifier *Kdm5d* and helicase *Helz2* (Fig S4E, Unstimulated). Four hours after TCR stimulation, *Gata3* expression was already higher in this group compared to wild-type controls (Fig S4E, 4 hours). By 24 hours, other Th2-specific genes such as *Ccr4* and *Gfi1* were significantly expressed in the Gata3^hi^ group (Fig S4E, 24 hours). Next, we conducted gene set enrichment analysis of the DEGs in Gata3^hi^ and Gata3^lo^ groups using the MSigDB Hallmark gene set collection^33^. The “IL2-STAT5 Signaling” module was enriched among upregulated genes at all time points in the Gata3^hi^ samples, but not Gata3^lo^ samples (Fig 4B). Gene sets associated with cell division and metabolism were also selectively upregulated in Gata3^hi^ samples after TCR stimulation, but not in the Gata3^lo^ samples, indicating that these samples were more responsive to TCR activation (Fig 4B). The IFN-γ module was also enriched in unstimulated after 48h of stimulation in Gata3^hi^ samples (Fig 4B). While IFN-γ signaling is commonly associated with Th1 effector function^7^, and anti-IFN-γ antibodies are typically used to reinforce Th2-polarizing conditions *in vitro*^37^, earlier studies from William Paul’s group suggested that the priming of Th2 populations, particularly the production of IL-4, could be initiated by early endogenous IFN-γ production and signaling^38,39^. Together, these data suggest that in the absence of *Tet2*, a pro-Th2 program is already established in naïve T cells and is further reinforced overtime upon TCR stimulation through differential activation of proliferative and cytokine signaling pathways and IL-4 production.

**Figure 4.**
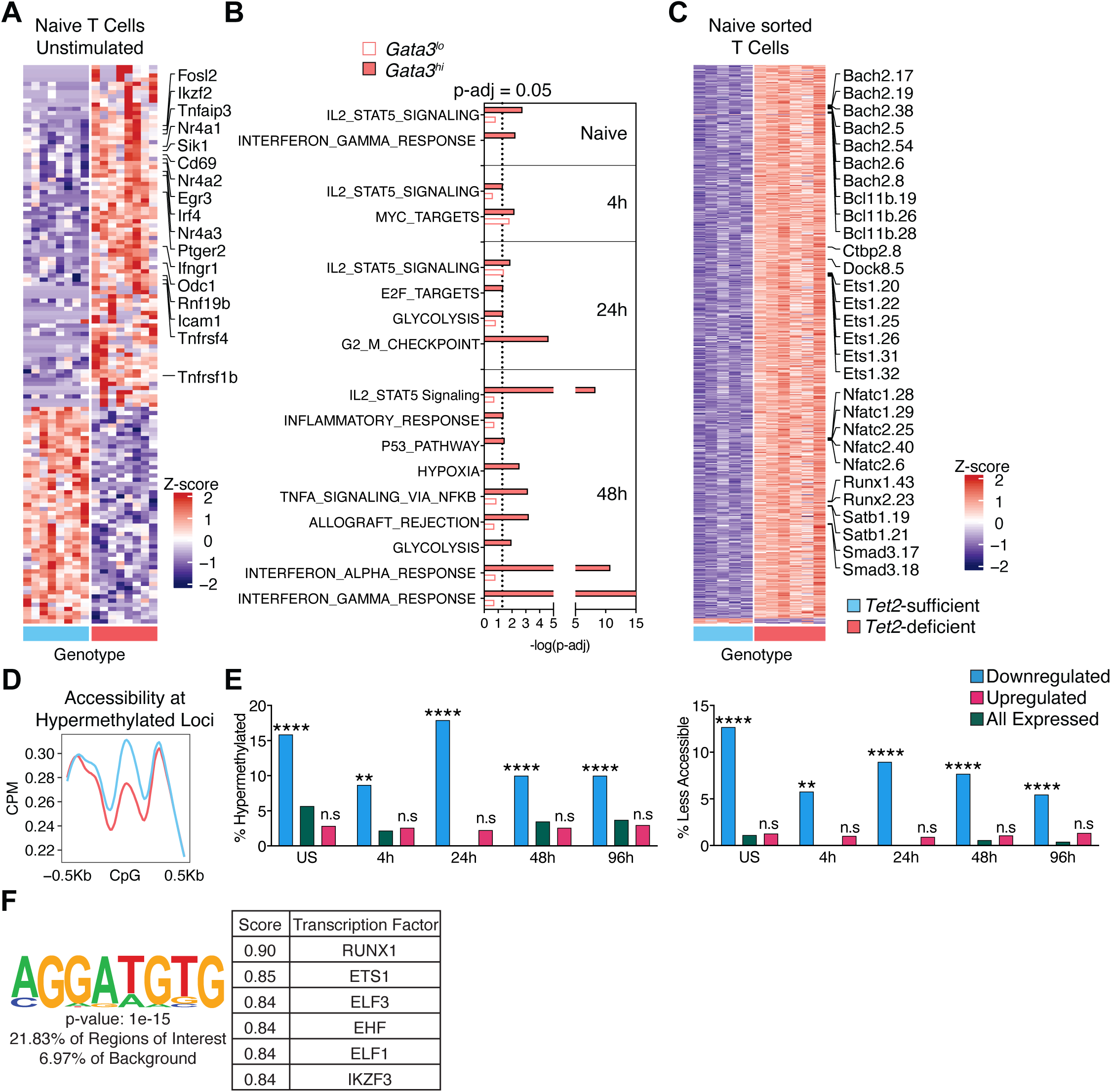
Epigenetic re-programming enables IL-4 production in the absence of *Tet2*. A, Z-score of normalized counts of differentially expressed genes in naïve *Tet2*-deficient and *Tet2*-sufficient CD4 T cells B, Enriched Hallmark gene sets in Gata3Hi *Tet2*-deficient relative to *Tet2*-sufficient samples at each culture time point. Also plotted are the p-adjusted values for Gata3Lo *Tet2*-deficient samples for each of these gene sets. C, Z-scored intensity of differentially methylated loci across naive CD4 cells from CD4-cre Tet2 fl/fl mice and littermates. D, Normalized accessibility derived from ATAC-seq of CD4 cells from CD4-cre Tet2 fl/fl mice and littermates, windowed around CpG loci that were hypermethylated in naïve CD4 cells from CD4-cre Tet2 fl/fl mice. E, Proportion of significantly down-regulated, up-regulated or total expressed genes with hypermethylated loci or decreased accessibility from Tet2-deficient CD4 T cells that were unstimulated or TCR stimulated for 96h. P-values represent Fisher’s exact tests for enrichment of epigenetic changes among these sets of genes. F, Consensus de novo motif from the sequences of regions less accessible in Tet2-deficient naive CD4 T cells along with p-value representing its enrichment and similarity score to known motifs identified from CHIP-seq of the indicated transcription factors that were most similar the identified de novo motif.

Our *in vitro* differentiation and sequencing data thus far were performed using naïve CD4 T cells from *Tet2*-/- and *Vav*-cre+ *Tet2*^fl/fl^ mice, along with their respective littermate controls, within our *Tritrichomonas*-colonized colony. The isolation of naïve CD4 T cells from these mice enabled a reductionist culture system, allowing us to identify an intrinsic bias towards Th2 differentiation under non-polarizing conditions. However, this approach does not completely exclude the effects of non-CD4 immune populations or systemic modulation by *Tritrichomonas*. To further characterize the role of the demethylase *Tet2* in establishing this intrinsic bias, we chose to study naïve CD4 T cells from CD4-cre *Tet2*^fl/fl^ mice that had not been exposed to *Tritrichomonas.* We first confirmed that under these conditions naïve T cells were also able to differentiate into Th2 cells under nonpolarizing conditions (Fig S4F), demonstrating that in the absence of *Tet2,* naïve T cells were prone to differentiate into Th2 cells independent of external environmental triggers. To directly interrogate the demethylase function of Tet2, we isolated naïve CD4 T cells from *CD4*-cre *Tet2^fl/fl^*mice and littermates and performed array-based methylation profiling as well as an assay for transposable-accessible chromatin sequencing (ATAC-Seq) from the same biological replicates. By comparing the log-transformed fraction of methylated to unmethylated DNA at a probe site between *Tet2*-deficient and wildtype samples, we obtained a set of differentially methylated CpG loci (DML) from the probe-based methylation data. As expected, most of the differentially methylated regions (DMLs) were hypermethylated in *Tet2*-deficient cells. Notably, several negative regulators of Th2 differentiation (Nfatc1^40^, Nfatc2^41^, Bcl11b^42^, Runx1^43^, Il6st^44^, Dock8^45^ and Lef1^46^) and IL4 production (Bach2^47^, Ctbp2^48^, Bcl11b^42^, Ets1^49^ and Lef1^50^) had sites with increased methylation in *Tet2-*deficient naïve T cells, which is in line with the increased IL-4 production observed upon TCR stimulation (Fig 4C).

We next compared our methylation data with the previously generated gene expression data. Overall, genes with hypermethylated loci in the *Tet2-*deficient samples exhibited lower expression levels, as indicated by log2 fold change (Fig. S4G). Importantly, this difference, which was already statistically significant in unstimulated naïve samples, became even more pronounced 96 hours after stimulation (Fig. S4G), at which point *Tet2*-deficient cells had adopted a prototypic Th2 identity (Fig. S4A-B). Furthermore, differences in methylation coincided with decreased chromatin accessibility at DMLs (Fig 4D), establishing a link between the demethylase function of *Tet2* and changes in genomic architecture at the hypermethylated sites. These observations suggest that *Tet2*-deficiency perturbs the epigenetic state of naïve CD4 T cells.

We next performed an integrated analysis of changes in both methylation and chromatin accessibility alongside with changes in gene expression in *Tet2*-deficient CD4 T cells at different timepoints of stimulation in non-polarizing cultures. Within down-regulated genes in naïve unstimulated *Tet2*-deficent cells, there was a significant enrichment hypermethylated loci or less accessible regions when compared to the presence of these loci and regions among all expressed genes (Fig 4E). No such enrichment was seen among the up-regulated genes (Fig 4E). We found similar enrichment in down-regulated but not up-regulated genes at 4h, 24h, 48h and even 96h after TCR stimulation (4h-96h, Fig 4E). These data suggest that *Tet2*-deficient naïve T cells are epigenetically poised to have reduced expression of certain genes throughout TCR-mediated differentiation into their effector state. These genes with reduced expression specifically included T cell signaling (Ubash3b^51^: US, 24h, 48h, 96h; Cers6^52^: 24h, Atp1b1^53^: US, 24h, 48h, Igsf23^54^: all timepoints) and modulators of cell activation and quiescence (Cd101^55,56^: US, 48h, 96h, Fam105a^57^, Lrig1^58^: 4h, 24h, 48h, 96h), further suggesting epigenetic determinants in *Tet2*-deficient T cells influence Th2 differentiation under non-polarizing conditions.

To probe the transcription factors associated with this epigenetic rewiring, we next performed transcription factor binding site motif analysis on the nucleotide sequences from regions that were less accessible in *Tet2*-deficient naïve CD4 cells. We found enrichment of motifs known to bind to several ETS family transcription factors (Fig S4H) and a de novo motif analysis specifically identified a motif with high similarity to those occupied by several validated negative regulators of Th2 differentiation, including those previously identified with hypermethylated sites, Runx1^43^ and Ets1^49^ (Fig 4F). Interestingly, Runx1 has been previously shown to interact with Tet2 in mediating demethylation of target genes^59^. Hypermethylation of loci associated with these critical negative regulators of Th2 differentiation (Fig 4F) along with decreased accessibility of their putative binding sites in Tet2-deficient naïve T cells indicates the demethylase function of Tet2 may help maintain the plasticity of naïve T cell differentiation by maintaining the expression and function of regulators of T helper differentiation. This plasticity is reproducibly lost in *Tet2*-deficient naïve CD4 T cells, resulting in increased autocrine IL4 production, collectively representing a loss of a critical checkpoint modulating Th2 differentiation.

## Dysregulated Th2 immunity predisposes allergic pathology

Given exogenous IL-4 was only sufficient to induce Th2 immunity in wildtype mice upon *Tritrichomonas* colonization (Fig 3I), we hypothesized, that in addition TCR stimulation and IL-4, other *in vivo* signals uniquely associated with *Tritrichomonas* colonization might play a role in inducing and maintaining the commensal-induced Th2 population in the gut. Previous work by Garrett, Locksley, von Moltke and colleagues has shown that IL-25 upregulation in response to *Tritrichomonas* was a key factor in establishing the type-2 innate immune circuit^14–16^. We had already observed significant correlation between tissue expression of *Il25* and the presence of Th2 cells in *Tet2*-deficient mice colonized with *Tritrichomonas* (Fig 2E), and thus we posited that IL-25 may also be required for the establishment and/or maintenance of Th2 immunity in response to Tritrichomonas colonization. We first analyzed the expression of the IL-25 receptor subunit IL-17RB on the surface of different intestinal lymphocytes (Fig 5A). Interestingly, Th2 cells but no other helper subsets in the intestines of *Tet2*-deficient mice expressed IL-17RB at levels to ILC2s, as measured by flow cytometry. Of note, absence of *Tet2* did not impact the level of IL-17RB expression in ILC2s (Fig 5A). We next tested the impact of IL-25 neutralization after chronic *Tritrichomonas* colonization in *Tet2*-deficient mice found that the Th2 population, normally induced by this commensal protist, was significantly reduced (Fig 5B). These data suggest that inducing persistent intestinal Th2 immunity to a commensal microbe like *Tritrichomonas* requires the microbe to induce a specific tissue signal, which indicates a change in the intestinal environment and the establishment of a specific immune response. This finding echoes studies showing that establishing long-lived Th2 cells in the lungs after type 2 immune challenges depends on the stress-induced cytokine IL-33^60^.

**Figure 5:**
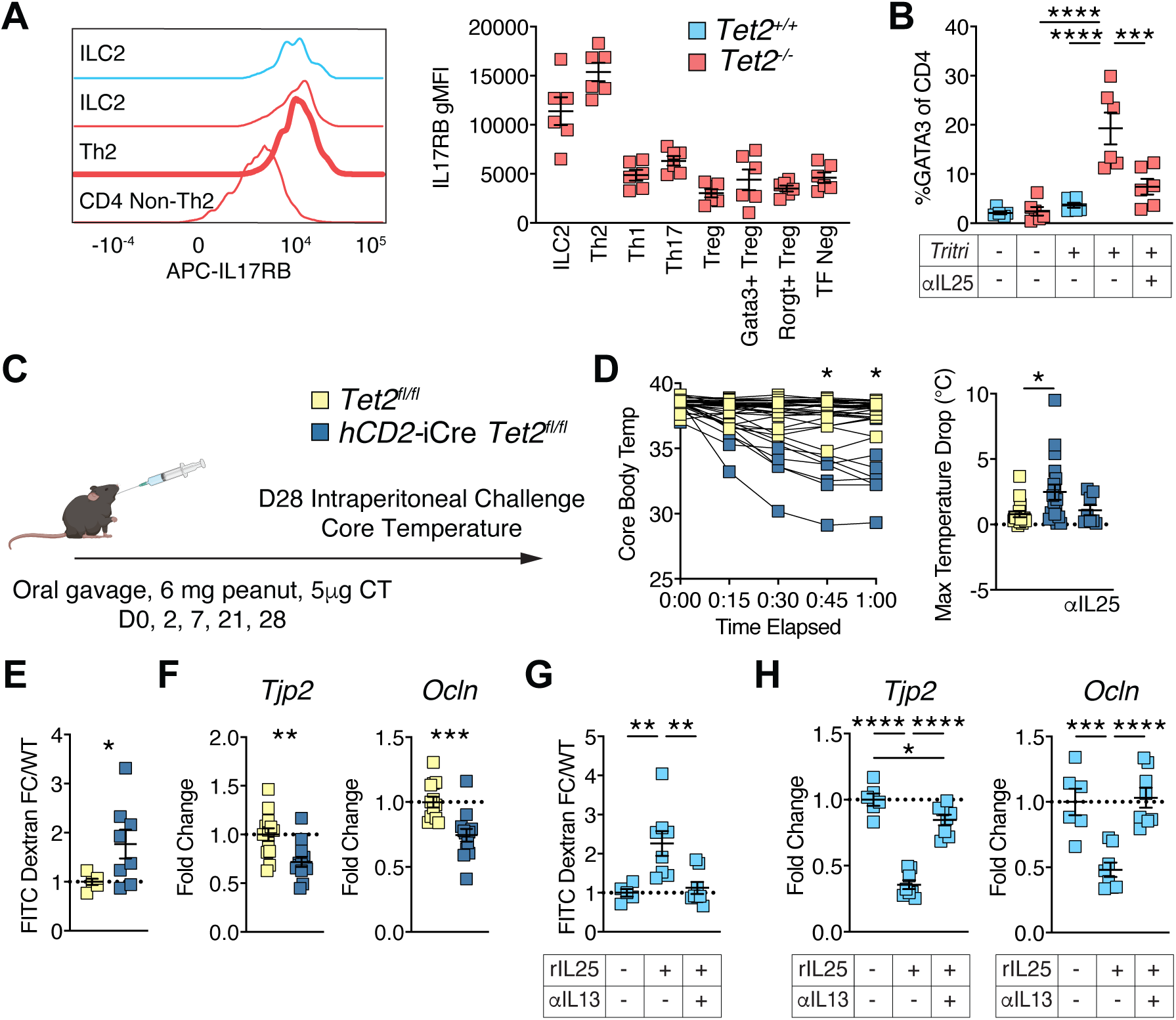
Tet2-deficiency predisposes mice to allergic pathology and leads to chronic IL25-dependent barrier dysfunction. A, Histogram (left) of IL17RB expression on indicated cell types from the jejunal lamina propria of *Tritrichomonas*-colonized wildtype (ILC2) and *Tet2*-deficient (ILC2, Th2, non-Th2) mice. Geometric MFI (right) of IL17RB expression across ILC2s and various CD4 T effector types from *Tet2*-deficient mice colonized with *Tritrichomonas*. B, Proportion of FOXP3-GATA3+ of CD4 T cells from the jejunal lamina propria of chronically *Tritrichomonas*-colonized and uncolonized *Tet2*-sufficient and *Tet2*-deficient injected every 3 days with anti-IL25 or isotype as indicated for 2 weeks. C, Sensitization and challenge schematic for the cholera toxin-peanut model of anaphylaxis. D, Serial temperature measurements (left) of hCD2-Cre Tet2fl/fl mice and littermates after I.P challenge with peanut allergen and maximum temperature drop (right) of untreated or anti-IL25 treated (during sensitization) hCD2-Cre Tet2fl/fl mice and littermates after peanut challenge. E, FITC-dextran permeability measured in the peripheral blood 3h after gavage of 0.1mg/g FITC-dextran. Represented as fold-change over littermate Tet2fl/fl. F, Quantitative PCR of *Tjp2* and *Ocln* measured relative to *Gapdh* and represented as fold-change over littermate Tet2fl/fl. G, FITC-dextran permeability for mice treated with vehicle, recombinant IL25 or recombinant IL25 and anti-IL13 was measured as before and represented as fold-change over vehicle treated wildtype mice. H, Quantitative PCR of *Tjp2* and *Ocln* relative to *Gapdh* in the jejunum of mice in (G) represented as fold-change over vehicle treated wildtype mice. Statistical analysis was performed by two-sided Student’s t-test in (D, E, F), and one-way ANOVA with Tukey multiple-comparison test in (B, D, G, H). p-values: * (<0.05) **(<0.01) *** (<0.001) **** (<0.0001). All panels representative of at least 2 independent experiments.

Persistent IL-25 signaling leads to chronic compositional and physical remodeling of the small intestine, including changes in tuft cells and intestinal length (Fig 1B, S1B). Previous studies using IL-25 transgenic mice have shown that IL-25 overexpression can increase susceptibility to allergic disease^61^. However, no studies have demonstrated that IL-25 induction in response to *Tritrichomonas* is associated with an increased propensity to develop allergic responses, nor have they identified potential mechanisms explaining why IL-25 overexpression is linked to an increased risk of allergy. To test the hypothesis that the induction of a chronic IL-25 signaling circuit associated with Th2 immunity in response to the commensal *Tritrichomonas* (Fig. 1H, Fig. S2C) increases susceptibility to food allergy, we utilized a peanut allergy model adjuvanted with a relatively low dose of cholera toxin (CT). This dose was generally insufficient to induce anaphylaxis in wild-type Tritrichomonas-colonized mice (Fig. 5C-D). In line with our hypothesis, *Tritrichomonas*-colonized hCD2-Cre *Tet2^fl/fl^* mice exhibited a significant increase in anaphylaxis compared to littermate control mice, as evidenced by a drop in core body temperature following systemic challenge (Fig 5C-D). Peanut antigen-specific IgE and IgG1 antibodies were equivalently induced in both genotypes, which is consistent with previous observations showing that the presence of antigen-specific antibodies does not correlate with allergic pathology in this model^62^ (Fig S5A). Furthermore, and as anticipated, hCD2-Cre *Tet2^fl/fl^* mice treated with an IL-25 neutralizing antibody throughout the sensitization period, no longer exhibited robust anaphylactic reactions upon peanut challenge (Fig 5D, right). These data indicate that chronic IL-25 signaling, resulting from persistent Th2 immunity in response to a commensal protist, has the potential to drive allergic immunopathology.

Building on our prior discovery of microbiota-dependent barrier dysfunction in *Tet2*^-/-^ mice^15^ and considering the reported link between barrier dysfunction and allergic responses^26,63^, we investigated whether persistent IL-25 signaling pathway, stemming from *Tritrichomonas* colonization in the context of *Tet2*-deficiency, promoted barrier dysfunction. In line with this hypothesis, hCD2-Cre *Tet2^fl/fl^* mice, which display persistent IL-25 signaling (Fig S3A) and allergic susceptibility (Fig 5C,D), exhibited heightened intestinal permeability, as evidenced by FITC-dextran assay (Fig. 5E), along with alterations in the expression of genes associated with barrier function^64^ (Fig. 5F). Interestingly, wild-type mice also developed barrier dysfunction upon *Tritrichomonas* colonization (Fig. S5B), which coincided with the acute increase in the IL-25 circuit (Fig 1h). However, consistent with the establishment of a chronic IL-25 circuit in *Tet2*-deficient mice, persistent barrier dysfunction was only observed in *Tet2*-deficient mice (Fig S5C, Fig 1H).

The compromise of intestinal barrier integrity has been implicated in various pathological processes, but identifying the specific signals responsible for barrier dysfunction has proven challenging in complex inflammatory environments ^63^. While previous studies have alluded to the role of type 2 inflammation in barrier dysfunction, our findings are unique in that they are measured in response to a commensal protist rather than pathogenic^65–67^ or chemical induced changes^68^ to the intestinal environment and do not rely on *in vitro* measurements. This is an important distinction as the presence of these pathways in homeostatic host-commensal interactions may represent a way to modulate allergic disease. To directly demonstrate whether activation of the IL-25 circuit could induce intestinal barrier dysfunction, we administered recombinant IL-25 to *Tet2*-sufficient mice in the absence of *Tritrichomonas* colonization. This was performed both in the presence and absence of a neutralizing α-IL13 antibody, which is essential for the propagation of the type-2 IL-25-IL13 circuit^19^. As anticipated, IL-25 injection led to an expansion of tuft cells in the epithelium in an IL-13-dependent manner (Fig S5D). IL-25 injection also disrupted intestinal barrier integrity, as evidenced by the FITC-dextran assay (Fig. 5G) and the downregulation of several genes associated with barrier function (Fig. 5H). Furthermore, IL-25-induced barrier dysfunction was contingent on IL-13 (Fig 5G-H). Taken together these results demonstrate that the engagement of the type 2 IL-25-IL-13 cytokine circuit, traditionally associated with establishing commensalism^14^ or clearing helminths^19,69^, can promote barrier dysfunction and allergic immunopathology when co-opted by aberrant and chronic adaptive responses (Fig S5E).

## Discussion

The mechanisms underlying the dysregulation of type 2 immunity in the intestinal environment and the relationship with food allergies remain poorly understood. Our study emphasizes that a combination of events, including the loss of a checkpoint allowing naïve T cells to produce IL-4 upon TCR stimulation and the presence of a microbe capable of inducing both T cell activation and tissue stress signals such as IL-25, predisposes individuals to chronic Th2 immunity and food allergy. Strikingly, in the absence of *Tritrichomonas* no Th2 immunity develops in our SPF *Tet2*-deficient mouse colony. Conversely, *Tritrichomonas* is sufficient to promote a chronic Th2 response. This unique feature of *Tritrichomonas* relies on its capacity to induce IL-25 and establish chronic commensalism even after the Th2 immune circuit is induced. The reasons why neither the transient ILC2-IL-25-IL-13 nor the persistent Th2-IL-25-IL-13 circuits succeed in clearing *Tritrichomonas*, and whether chronic colonization of the intestine by *Tritrichomonas* in the absence of a persistent IL-25-IL-13 circuit may be evolutionarily beneficial, remain to be determined.

Unlike Th17 and Th1 immunity, adaptive Th2 immunity is typically excluded from the small intestine. This is evident from the fact that a commensal protist like *Tritrichomonas* induces only a transient innate ILC2 circuit in wild-type mice^14^. The regulation of the type 2 intestinal circuit thus has been described at the level of innate lymphocytes, where the expression of the transcription factor A20 is necessary to restrain the activation and function of ILC2s in the presence of *Tritrichomonas*^16^. Dysregulation of ILC2 activation was shown to lead to lower body weight and a chronically lengthened small intestine, presumably a metabolic adaptation to a chronic type 2 circuit^16^. Our findings revealing that chronic Th2 immunity weakens barrier function and increases susceptibility to immunopathology, further underscore the harmful consequences of sustained Th2 immunity in the gut. IL-4 plays a central role in Th2 immunity, serves as a crucial factor for the differentiation of Th2 cells and functions as a key type-2 effector cytokine.^5^ It regulates both innate and adaptive immune responses, as well as tissue function and remodeling^5^. The dual nature of IL-4, a driver and effector cytokine of Th2 immunity, highlights the critical need to regulate its production in naïve T cells to prevent dysregulated type-2 immune responses. Our findings highlight the role of Tet2 in controlling IL-4 production by naïve T cells and their capacity to undergo Th2 polarization in the absence of exogenous IL-4, which is typically produced by innate cells such as basophils upon infection with pathogens^70,71^.

Our study also identifies IL-25 as a key trigger for the development of barrier dysfunction, providing a mechanistic basis for the previously described allergic predisposition observed in an IL25-transgenic mouse model using ovalbumin antigen adjuvanted with alum^61^. Overall, the mechanisms underlying the increased incidence of food allergy and the development of pathogenic Th2 responses are complex and likely involve both genetic and environmental factors^72^. Our findings suggest that a combination of a genetic defect (such as *Tet2* deficiency) and the presence of a commensal protist is necessary and sufficient for the development of pathological Th2 responses. Naïve T cells from the Th2 prone BALB/c mouse strain have a significantly higher propensity to produce IL-4 upon TCR stimulation than C57BL/6 naïve CD4 T cells^73,74^. Interestingly, BALB/c mice harbor an inactivating mutation in the *Tas1r3* gene that renders tuft cells incapable of recognizing *Tritrichomonas*-produced succinate and initiating the IL-25 circuit^75^. This raises the question of whether selection pressures for this trait in BALB/c mice may have been driven by immunopathology associated with chronic Th2 immunity under homeostatic conditions. Studying genetic variants in the context of immune biases and host-microbial interactions across laboratory mouse strains could therefore be informative in uncovering other mechanisms important for maintaining homeostasis at barrier surfaces.

Whether a commensal triggered Th2 response and/or genetic driven alterations in *Tet2* expression are involved in human allergic responses, such as drug hypersensitivity responses observed in patients with T-cell leukemias^76^, remains an intriguing possibility to be investigated. Indeed some evidence implicating TET2 function in T cells from allergic rhinitis patients has already emerged^77^. Furthermore, our study identifies changes at the level of methylation and genomic architecture that predispose Th2 differentiation, and potentially serves a predisposing factor for allergic responses. Changes in DNA methylation have been described in patients with allergic type diseases and represents interest in identifying epigenetic changes associated with these diseases and understanding the molecular pathways to which they may correspond^78–81^. While our study reveals a role for *Tet2*, multiple mechanisms likely regulate IL-4 production by naïve T cells upon TCR activation. We highlight a key mechanism and context where dysregulation can lead to immunopathology. Identifying T cell-intrinsic pathways for Th2 development is crucial for understanding and preventing the growing prevalence of allergic diseases.

## Author contributions

Conceptualization: KS, BJ, RH

Methodology: KS, BJ

Investigation: KS, MP, LC, SP, HA, RH, AT, JP

Visualization: KS

Funding acquisition: BJ, RH, MM

Project administration: BJ

Supervision: BJ, KS

Writing – original draft: KS, BJ

Writing – review & editing: RH, SR, MP, MM, LBB

## Acknowledgements

We would like to thank the support staff at the University of Chicago Animal Resources Center for making all our studies possible. Additionally, we would like to thank the Human Disease and Discovery Core, Functional Genomics Core, and the Flow Core at the University of Chicago. We would like to thank EK, AT and JE for technical help. We would like to thank LG for discussion and TM and LBB for critical reading and discussion of the manuscript.

## Methods

### Mice

Mice between 6 and 20 weeks of age were used as specified. Mice were housed in specific pathogen-free (*Helicobactor hepaticus* free, murine norovirus free) or germ-free conditions as specified. *Tet2*^-/-^ mice were obtained as previously described^26^. The following strains were purchased from Jackson lab and crossed in-house: *Tet2*^fl/fl^ (#017573), *Vav1*-Cre (#008610), hCD2-Cre (#008520), *Il5*-Cre (#030926), *Cd4*-Cre (#022071), *Rag2^-/-^* (#008449). Germ-free mice were generated previously described^26^ and housed and maintained by the University of Chicago gnotobiotic facility. All mice were maintained on a standard chow diet and husbandry and experiments were in accordance with policies and protocols approved by the University of Chicago Institutional Animal Care and Use Committee.

### Mouse treatments

Depleting or blocking antibody treatments were injected intraperitoneally (I.P) in 200uL at the indicated doses every 3 days for the duration of the experiment: 500ug anti-IL25 (Amgen, mouse IgG1 isotype), 200ug anti-IL13 (Janssen), 200ug anti-CD4 (Bioxcell, rat IgG2b isotype), 200ug anti-IL4 (Bioxcell, rat IgG1 isotype). For rIL25 treatment, 300ng of IL25 (R&D) was injected I.P for 3 days followed by end point analysis. For rIL4c treatment, 1ug of rmIL4 (R&D, CF) was incubated with 5ug of anti-IL4 (Bioxcell) for 30 minutes at room temperature and injected IP in 100uL final volume. For metronidazole treatment, mice were provided with 2.5g/L metronidazole (Sigma) and 1% sucrose ad libitum in drinking water. Bottles were replaced weekly.

### Tissue Processing

Mouse small intestines were removed and washed in cold PBS. 1-cm segments were cut from relevant segments for RNA extraction and histology readouts. 12-cm of jejunum was measured for lamina propria isolation^83^. Briefly, intestines were cut open longitudinally, washed in cold PBS to remove luminal contents and then were shaken 3 times in IEL medium (2mM EDTA in HBSS) at 37°C and 250rpm for 10 minutes each. Between shakes, the tissue pieces were washed with warm HBSS on 100um filters to aid in removing epithelial cells. After the IEL shakes, the tissue pieces were shaken in LPL medium (20% FBS, 0.05mg/mL DNAse I, 1mg/mL Collagenase A in RPMI) for 30 minutes. After the LPL shake, digested issue pieces were passed through a 100um filter, centrifuged, and stained for FACS as described below. Mesenteric lymph nodes and Peyer’s patches were isolated and separated by segment as previously described.^84^ Samples were collected in complete RMPI (10% FBS, 1% PenStrep Glutamine) and shaken in 1mL digestion medium (1mg/mL Collagenase VIII in complete RMPI) 30 minutes at 37°C at 250rpm. Digestion was halted by adding 10uL 0.5 M EDTA and placing samples on ice for 10 minutes. The media and remaining tissue were passed through 100uM filters and mashed. Digested and dissociated samples were washed once and then stained for FACS.

### Epithelial cell isolation

Epithelial cells from segments of jejunum were isolated as previously described^85^. Briefly 7cm of intestinal segment was excised and flushed with cold PBS. The distal end of the segment was tied to the end of a gavage needle using suture and the proximal end of the segment was inverted and tied off at the other end. The gavage needle was attached to a syringe. This inverted intestine ‘pouch’ was initially submersed in cold PBS for an additional wash and then placed in a glass tube containing BD Cell Recovery Solution (Corning, Cat. 354253). The intestinal pouch was inflated and incubated in the recovery solution for 5 minutes in the cold room. Every 5 minutes for 6-8 cycles, the intestinal pouch was deflated and re-inflated, allowing sheets of epithelium to detach. The glass tubes were then collected, their contents were spun down at 2000g for 10 minutes and then stained and processed for flow cytometry as mentioned below.

### Cell Staining and Flow Cytometry

Cells were stained with FC block (BD #553142, 10 min), fixable live dead dye (Invitrogen #L34957 or BioLegend #423105, 15 min), and surface markers (25 min). Cells were then fixed and stained intracellularly using the eBioscience Foxp3 kit (#00-5523-00) according to manufacturer instructions. Staining panels were generated using antibodies from the following:

CD45 (Pac Blue; Clone 30-F11; BioLegend 103126)

CD45 (AF532; Clone 30-F11; eBioscience 58-0451-82)

CD45.2 (BUV395; Clone 104; BD 553772)

CD45.1 (Pac Blue; Clone A20; BioLegend 110722)

TER119 (BV605; Clone TER-119; BioLegend 116239)

F4/80 (PEcy7; Clone BM8; BioLegend 123110)

Ly6G (BV750; Clone 1A8; BD 747072)

Epcam (BV605; Clone G8.8; BioLegend 118227)

Epcam (PerCP/Cy5.5; Clone G8.8; BioLegend 118220)

CD90.2 (BV421; Clone 53-2.1; BioLegend 140327)

CD90.2 (BV510; Clone 30-H12; BioLegend 105335)

CD3e (BUV737; Clone 145-2C11; BD 612771)

Tcrb (PE-CF594; Clone H57-597; BD 562841)

CD4 (BUV615; Clone GK1.5; BD 613006)

CD4 (FITC; Clone GK1.5; BioLegend 100406)

CD8α (BUV395; Clone 53-6.7; BD 563786)

CD8β (BV480; Clone H35-17.2; BD 746835)

TCRgd (BV650; Clone GL3; BD 563993)

GATA3 (PerCP-eflour710; Clone TWAJ; eBioscience 46-9966-42)

Ki67 (PE; Clone SolA15; eBioscience 12-5698-82)

KLRG1 (BV711; Clone 2F1; BD 564014)

IL17RB (APC; R&D FAB10402A)

Rorγt (BV786; Clone Q31037; BD 564723)

Foxp3 (eflour450; Clone FJK-16s; ThermoFisher 48-5773-82)

Tbet (PE; Clone 4B10; BioLegend 644810)

CD44 (PE-Cy7; Clone IM7; BioLegend 103030)

CD62L (PE; Clone MEL-14; BioLegend 104408)

Samples were acquired on a LSR Fortessa X20 (BD) or Cytek Aurora machine and data was analyzed using FlowJo (v10, TreeStar).

### Tritrichomonas identification, isolation, and colonization

Ceca from *Tritrichomonas*-colonized mice were excised, and their contents were removed by rinsing exposed contents in antibiotics-containing PBS (50ug/mL vancomycin (Sigma #V2002), 100mg/mL streptomycin (Sigma #S9137), 100U/mL penicillin (Sigma #P3032); ABX-PBS). This slurry was passed through a 100um filter and spun down at 1000rpm for 7 minutes after which it was re-suspended in 5mL 40% v/v percoll made in ABX-PBS and overlaid on 5mL 80% v/v Percoll (GE). The percoll gradient was spun for 15 minutes at 1000g with no brakes at room temperature. The interphase was collected and washed twice in fresh PBS before sorting 2e6 protists per mouse to be colonized. *Tritrichomonas* burden was measured by extracting DNA from cecal or colonic contents using the Qiagen Fast Stool kit (Qiagen). Contents were homogenized in 1mL InhibitEx buffer (Qiagen) in 2mL screw top tubes filled with 0.5mL of 0.1mm glass beads (Biospec) using the Omni Bead Ruptor Elite homogenizer prior to DNA extraction. PCR was performed as below using 10ng of DNA as starting material.

### RNA/DNA isolation and Quantitative RT-PCR

Tissues stored in RNAlater (Qiagen) for 24-48h at 4C were transferred to RLT+ containing 2-mercaptoethanol and homogenized using an equal mix of 0.5 mm and 1.0 mm zirconium oxide beads (Next Advance) and a bead homogenizer. Cells were stored directly in RLT+ containing 1% 2-mercaptoethanol and frozen at -80°C. RNA for all samples was extracted using the Qiagen RNeasy kit according to manufacturer instructions. For samples for transcriptional and genomic analysis RNA and DNA was extracted using the Qiagen All Prep RNA/DNA kits according to manufacture instructions. Reverse transcription with 500ng of total RNA was performed using a GoScript Reverse Transcriptase kit (Promega) and PCR was performed on a Roche Light Cycler 480 machine using SYBR Advantage qPCR Premix (Clontech). Parameters for amplification: denature for 10s at 95C, anneal for 10s at 60c and extension for 10s at 72C. Relative expression was calculated using 1000* 2^^-βCT^ with *Gapdh* as the housekeeping gene. To account for technical variation, expression was normalized to wildtype or wildtype uncolonized samples. The primer sequences are as follows:

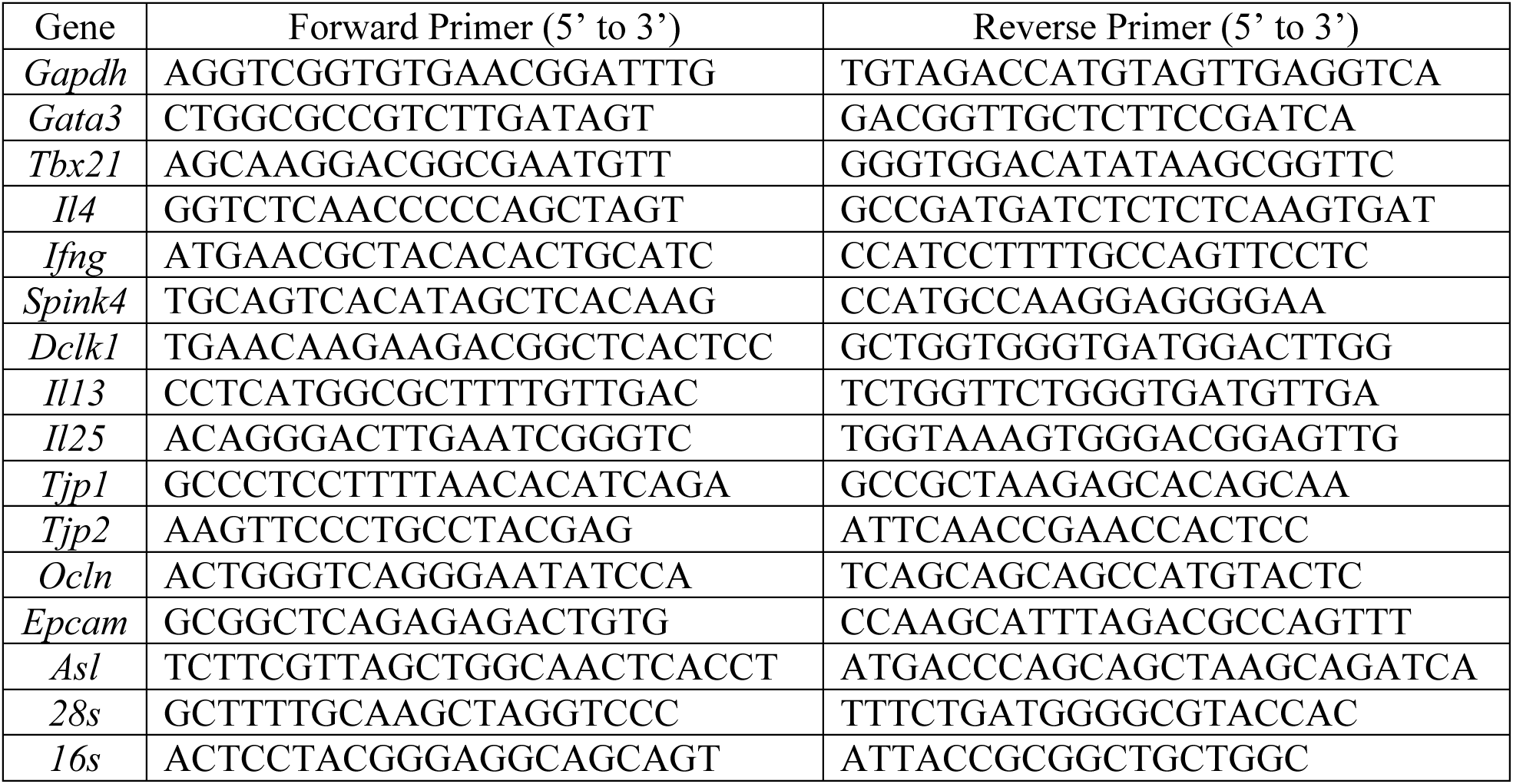

### Library preparation and sequencing

RNA quality and quantity were assessed using the Agilent Bio-Analyzer. Strand-specific RNA-SEQ libraries were prepared using a TruSEQ mRNA RNA-SEQ library protocol (Illumina provided). Library quality and quantity were assessed using the Agilent bio-analyzer, and libraries were sequenced using an Illumina NovaSEQ6000 (Illumina provided reagents and protocols).

DNA samples were quantitated by Qubit and bisulfite conversion was performed using a Zymo kit (D5004) and a Zymo provided protocol. Array hybridization and scanning were performed using Illumina Infinium Mouse Methylation BeadChips and the Illumina IScan using reagents and protocols provided by Illumina.

### ATAC-Seq Sample Preparation

ATAC-seq samples were prepared following an updated version of Omni-ATAC as described^86^. First, naive CD4+ T cells (Live CD4+CD25-CD62L+CD44lo) were sort-purified from Tet2 F/F (5 biological reps WT) and Tet2 F/F CD4-cre mice (6 biological reps Tet2 KO). 60,000 naive CD4+ T cells from each biological replicate were spun down for nuclei isolation followed by transposition using Tn5 transposase from the Tagment DNA (TD) TDE1 Enzyme and Buffer Kit (Illumina, cat. no. 20034197). After transposition, DNA Clean and Concentrator-5 Kit was used to purify the DNA fragments. Barcoding of the transposed fragments was performed with 5ul of IDT® for Illumina Nextera DNA Unique Dual Indexes Set A and library quantification was performed using NEBNext Library Quantification Kit to determine the number of additional PCR cycles for proper amplification of libraries after the initial 3 PCR cycles. For all samples, 6 additional cycles were performed for a total of 9 PCR cycles for library amplification. Samples were submitted to the University of Chicago Genomics Facility for 50bp paired-end sequencing on a NovaSeq X.

### Transcriptional Analysis and Visualization

Raw counts for bulk intestinal RNAseq data, processed as previously described^26^, were obtained from GSE99333. High quality raw reads (phred score >28) from *in vitro* T cell culture experiments were aligned using STAR (v2.6.1d, GRCm38, Gencode vM25) and summarized with featureCounts (subread v1.5.3). Subsequent analysis was performed in R (v4.0.3). Batch correction for experimental batches was performed using ComBATseq (sva v3.38.0), and differential expression analysis was performed using DESeq2 (v1.30.1).

Differentially expressed genes (adjusted P-value < 0.05 (Benjamini-Hochberg correction), |Log2FoldChange| > 0.25) at each timepoint were calculated from batch-corrected matrices using a negative binomial generalized linear model with design coefficients of genotype and/or expression of *Gata3*. These lists for each relevant comparison were tested for overrepresentation of pathways and gene sets using hypergeometric tests via the enricher() function in clusterProfiler. Annotated gene sets were derived from the molecular signatures database (MSigDB) and separate analyses were conducted for genes that were up-regulated and down-regulated (positive and negative log fold change, respectively) in our dataset to identify gene sets that were up-regulated and down-regulated as opposed to simply over-represented among differentially expressed genes. The background set of genes in these tests included all the genes represented in the unfiltered results table from DESeq2 and the threshold for adjusted p-values (Benjamini-Hochberg correction) of less than 0.05 determined set enrichment. Overrepresentation of pathways from publicly available a human naïve CD4 RNAseq dataset was similarly performed^87^. Briefly, differential expression of genes in CD4 T cells after activation was performed as above between individuals who had persistent allergy (as defined by original authors) and non-atopic controls. The upregulated and downregulated gene lists were then separately tested for overrepresentation of pathways and gene sets as above using the MSigDB.

For intestinal cell composition analysis from bulk RNAseq data, cell signatures were obtained from PanglaoDB^27^. For interrogating GATA3-regulated genes in non-polarizing cultures, we used gene lists from published ChIP-seq and RNA-seq of naïve and differentiated T helper cells ^32^. Using these curated gene lists, we inputted DESeq2 normalized count matrices, tested for enrichment of sets between *Tet2^+/+^* and *Tet2^-/-^* samples using gene set enrichment analysis (GSEA)^88^ to obtain normalized enrichment scores (NES) and false discovery rates (FDR) for each set, which were plotted with significant results highlighted in Figure 1 and S1.

Heatmaps were plotted using the ComplexHeatmap package (v2.6.2) in RStudio. A normalized counts matrix was obtained using the variance stabilizing transformation in DESeq2, and row-centered z scores were calculated for each gene.

### Epigenomic Analysis ATAC-Seq

ATAC-seq reads were preprocessed using the ‘atacseq’ pipeline (v2.1.2) from nf-core with the mouse genome build GRCm38^89^. Then, to perform principal component analysis (PCA) and to determine differentially accessible peaks, broad consensus peak counts for naïve samples were normalized using limma’s ‘voom’ function (v3.58.1)^90^ in R (v4.3.3) via RStudio (v 2024.04.2+764). PCA performed on the top 30% most variable peaks after normalization indicated that samples were separable along PC2 by sex, as well as separable along PC1 by *Tet2* genotype. Based on these results, sex was included as a separate additive term in the linear model that was chiefly used to test for differential accessibility of genomic regions based on genotype (∼ 0 + Tet2_genotype + sex). Specifically, differential accessibility testing was performed by fitting a genotype-specific contrast (CD4-cre *Tet2*^fl/fl^ compared to *Tet2*-sufficient littermates) using limma’s ‘contrasts.fit’ followed by empirical Bayes moderation of standard errors using limma’s ‘eBayes’. As part of the nf-core pipeline, peaks were annotated such that they were associated with the gene which had closest transcription start site to the peak.

### DNA methylation profiling by array

DNA methylation probe intensities were preprocessed using the ‘openSesamè function from the SeSAme package (v1.20.0)^91^ in R (v4.3.3) with the settings recommended for the Infinium Mouse Methylation BeadChip (referred to in SeSAme as MM285). Next, M-values (the log of the ratio of methylated versus unmethylated intensities of a probe) were calculated for each probe. PCA was then performed on the top 30% most variable M-values for naïve samples. Samples separated along PC1 by sex, which explained 55% of the variance, in addition to separating along PC3 (6% variance) by *Tet2* genotype. Due to this, as in the ATAC-seq analysis, sex was included in linear modeling to determine differentially methylated loci. This linear modeling was performed via limma’s ‘contrasts.fit’ and ‘eBayes’ functions, as in the ATAC-seq analysis. M values, as opposed to beta values (the fraction of the methylated probe’s intensity compared to total intensity), were used for the modelling, as M values have been shown to have a better accuracy for such analysis^92^. Probes were labeled with a unique identifier including a gene name if they were in a likely promoter region for that gene, defined by being within 1500 base pairs (bp) of the transcription start site, or if they were within an exon or intron of the gene. The heatmap of differentially methylated loci (Fig 4C) was plotted using the ComplexHeatmap package (v2.6.2) in RStudio, as above.

Accessibility at differentially methylated CpG loci was visualized using merged bigWig files. To generate bigWig files, aligned bed files were merged and converted to bigWig using bedtools (v2.3.1). Merged bigWig files and bed files containing the coordinates of CpG loci of interest were used to generate matrices containing normalized ATACseq counts surrounding these loci with deepTools (v3.5.5). These matrices were then imported into R Studio and plotted with ggplot2.

## Helminth propagation and infection

*Strongyloides venezuelensis* was generously provided by Daria Esterhazy (University of Chicago). To propagate and maintain cultures, NSG mice (Jax #00557) were infected subcutaneously with 10,000 L3 larvae. Fecal pellets from infected mice were homogenized in water and spread onto filter paper that was partially submerged in tap water in a beaker. The beaker was loosely covered with plastic film and placed at 30°C for 4 days after which water was collected and fresh larvae were allowed to settle at room temperature. Larvae were counted and resuspended in tap water for an infection dose of 700-1000 L3 in 100uL. Experimental mice were infected and sacrificed at day 7 to assess priming of intestinal Th2 responses in mesenteric lymph nodes.

## Naïve CD4 cultures

Naïve CD4 T cells were isolated from spleens of 6-8 week *Tet2^-/-^* or *Tet2^+/+^* mice by homogenizing spleens in RBC lysis (R&D #WL2000) and then enriching for naïve cells using MACS enrichment (Miltenyi #130-104-453). Cells were then further sort purified (CD4+ CD8-CD62L+ CD44-) and seeded at 1e5 cells/well in 96 well plates pre-coated overnight with 5ug/mL anti-CD3 (Biolegend #100302). Cells were cultured in complete RPMI (10% FBS, 1% PenStrep Glutamine) with soluble anti-CD28 (Biolegend #102102) and 10ng/mL mIL2 (Miltenyi #130-107-760) for non-polarizing cultures. For Th2 polarizing cultures, 10ng/mL IL-4 (Miltenyi #130-107-760) was added and for Th2-blocking cultures, 10ug/mL anti-IL-4 (Bioxcell #BE0045) was added. Cells were cultured for 48 hours and then split and reseeded in the same media as secondary cultures and analyzed at d4-6. For RNA, cells were collected in RLT+ and immediately stored at -80°C. Cell supernatants were frozen at -20°C before analysis of cytokines by ELISA.

## *In vivo* intestinal permeability

*In vivo* intestinal permeability was assessed by 4kDa FITC-dextran (Sigma-Aldrich #46944). Mice were deprived of food and water for 5 hours and then orally gavaged with 60mg/kg of FITC-dextran in PBS. Two hours later, venous blood form the maxillary sinus was collected, and spun down for 10 minutes at 10,000rpm. The supernatant was collected and measured for FITC-dextran using an excitation of 490nm and emission of 520nm on a fluorescent plate reader.

## Cholera toxin sensitization and peanut challenge

Crude peanut extract (CPE) was prepared from roasted unsalted peanuts. Briefly, peanuts were ground to and added to 20mM Tris buffer (pH 7.2) at 25 grams peanuts per 20mL 20mM Tris. The solution was stirred at room temperature for 2 hours and then centrifuged at 3000 g for 30 minutes. The aqueous fraction below the upper most fat layer was carefully removed and measured by BCA assay for protein content. Mice were sensitized with 6mg of peanut extract and 5ug of cholera toxin in 200uL by oral gavage on day 0, 2, 7, 14 and 21. On day 28, mice were challenged I.P. with 1mg of peanut extract in 200uL. Body temperature was measured using a rectal probe prior to challenge and every 15 minutes after challenging up to one hour. Mice were bled on day 14 and day 27 to assess antibody responses in the serum and were bled after challenge on day 28 to assess systemic mast cell degranulation.

## Enyme-linked immunoassay

Commercial ELISA kits were used to measure IL4 production and IFNγ according to manufacturer instructions (Invitrogen). Peanut specific IgE and IgG1 was measured by first coating high-binding 96-well plates (Corning 3690) with 20μg/mL CPE in 100mM Na_2_CO_3_ overnight at 4C.Plates were washed three times in PBS plus 0.05% Tween-20 (PBS-T) and blocked with 150μL 1% BSA in PBS-T for 2 hours at RT and then washed 1 time with PBS-T. Mouse serum obtained via submandibular bleed was diluted in blocking buffer and added as 25μL per well and incubated for 1 hour at RT followed by 3 washes. 50 μL horseradish peroxidase-conjugated anti-mouse IgE or IgG1 (Southern Biotech) diluted 1:1000 in blocking buffer was added to well and incubated for 45 minutes at RT and then washed 5 times. 50μL TMB substrate was added, and the reaction was stopped by adding 50μL 1N H_2_SO. Absorbance at 450nm was read immediately after stopping the reaction.

---

**Supplemental Figure 1, related to Figure 1:**
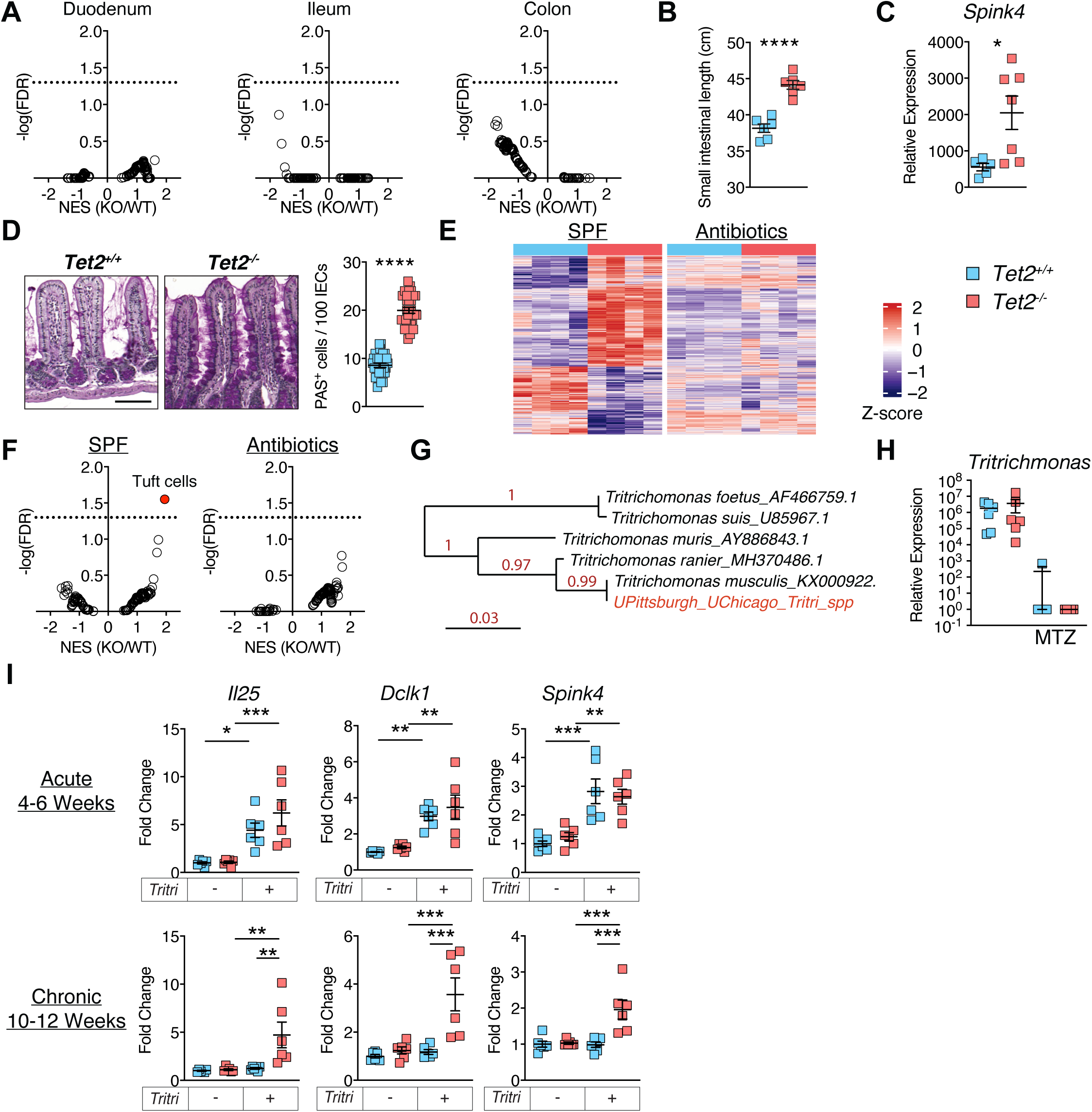
A, Volcano plot of normalized enrichment scores and adjusted p-values for various cell signatures from normalized RNAseq expression data from Tet2+/+ and Tet2-/- duodenum, ileum and colon tissue. B, Small intestinal length from Tet2+/+ and Tet2-/- mice C, Quantitative PCR of jejunal segments from SPF and GF Tet2+/+ and Tet2-/- mice expressed relative to *Gapdh*. D, Formalin fixed segments of jejunum were embedded in paraffin and subjected to periodic acid-Schiff staining. PAS+ cells were quantified in the epithelium as a proportion of total epithelial cells. E, Normalized counts of genes that were differentially expressed between SPF Tet2+/+ and Tet2-/- jejunum were Z-scored across all conditions and plotted for all conditions. F, Volcano plots of normalized enrichment scores and adjusted p-values for cell signatures from normalized RNAseq expression data from the jejunum of sham or broad-spectrum antibiotics-treated Tet2+/+ and Tet2-/- mice. G, Phylogenetic comparison of the amplified 28s ITS sequence from the microbiota of SPF mice in our facility with published 28s ITS sequences. H, Quantitative PCR of Tritrichomonas relative to 16s in cecal contents from Tet2+/+ and Tet2-/- mice treated with metronidazole or control water for 4 weeks I, Quantitative PCR of IL-25 circuit genes relative to Gapdh in the jejunum of acutely and chronically colonized Tet2+/+ and Tet2-/- mice. Represented as fold change over uncolonized littermate Tet2+/+ mice. Statistical analysis was performed by two-sided Student’s t-test in (B,C,D) one-way ANOVA in (I) with Tukey multiple-comparison test. p-values: * (<0.05) **(<0.01) *** (<0.001) **** (<0.0001). Panels b-d and g-i are representative of at least 2 independent experiments.

**Supplemental Figure 2, related to Figure 2:**
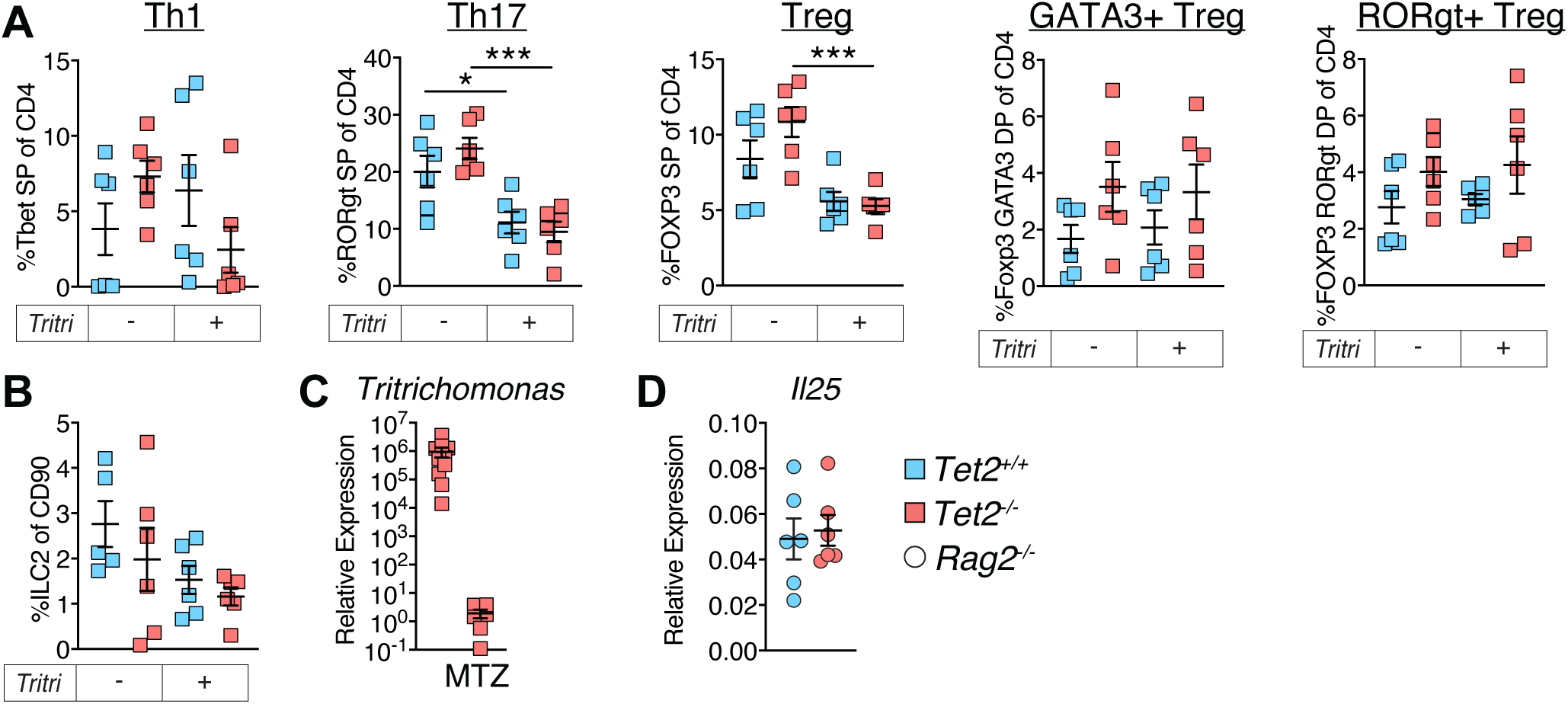
A, Proportion of CD4 effector types from the jejunal lamina propria of uncolonized and 4 week Tritrichomonas-colonized Tet2+/+ and Tet2-/- mice. B, Proportion of ILC2s of uncolonized and Tritrichomonas-colonized Tet2+/+ and Tet2-/- mice. C, Quantitative PCR of Tritrichomonas relative to 16s in cecal contents from Tet2+/+ and Tet2-/- mice treated with metronidazole or control water for 4 weeks D, Quantitative PCR of *Il25* relative to *Gapdh* in the jejunum of Rag2-/- Tet2+/+ and Rag2-/- Tet2-/- mice Statistical analysis was performed by one-way ANOVA in (A) with Tukey multiple-comparisons test. p-values: * (<0.05) **(<0.01) *** (<0.001) **** (<0.0001). All panels representative of at least 2 independent experiments.

**Supplemental Figure 3, related to Figure 3:**
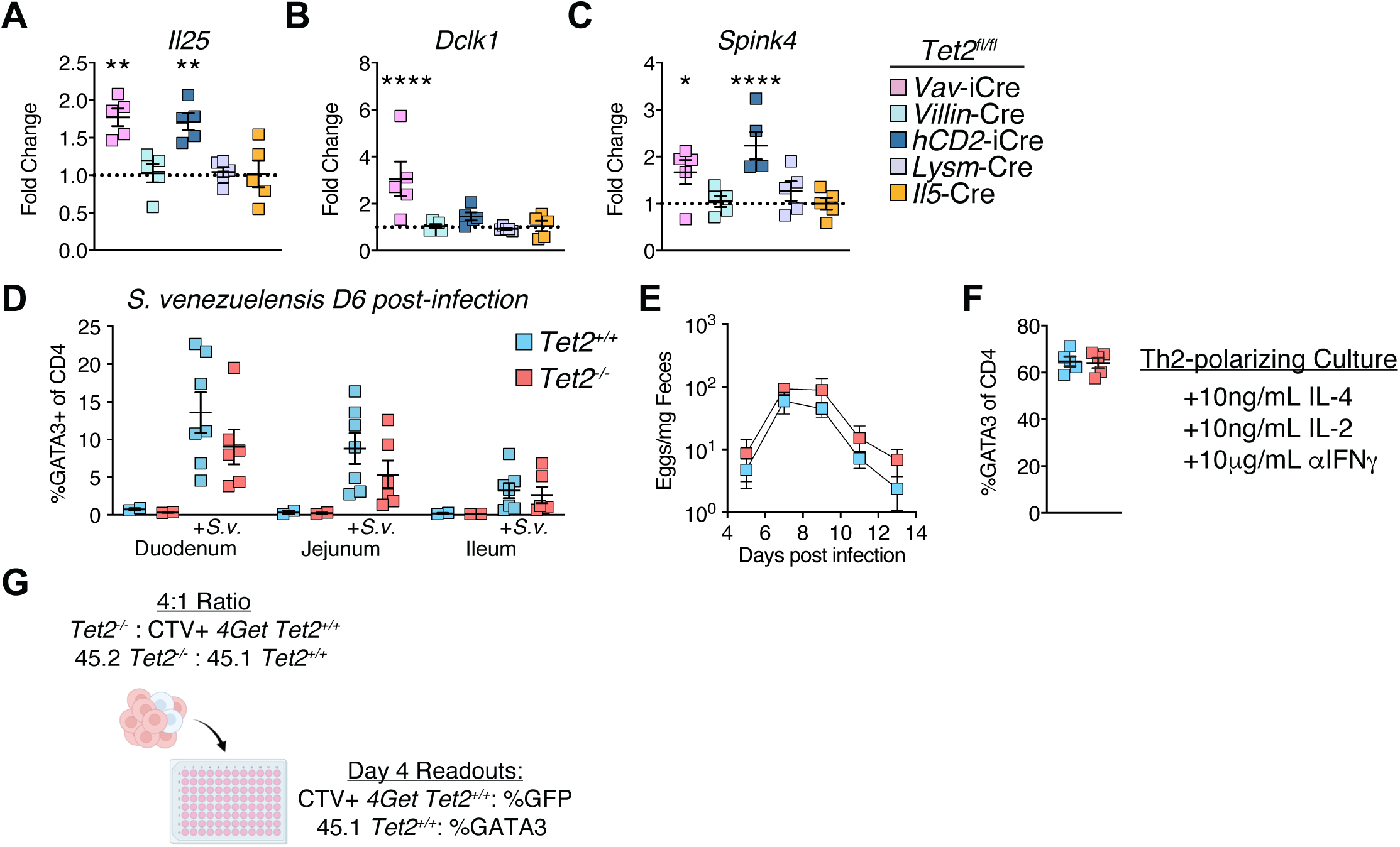
A, B, C, Quantitative PCR of Il25 (A), Dlck1 (B) and Spink4 (C) relative to Gapdh in the jejunum of Tet2 fl/fl mice with various Cre drivers. Expressed as fold change of Cre-expressing lines over littermate Tet2 fl/fl mice D, GATA3+ Foxp3-proportion CD4 T cells in mesenteric lymph nodes draining indicated intestinal segments in Tet2+/+ or Tet2-/- mice 6 days after being infected with 1000 S. venezuelensis L3 larvae subcutaneously. E, S. venezuelensis eggs quantified in fecal pellets from infected Tet2+/+ or Tet2-/- mice over 2 weeks of infection. F, Proportion of Foxp3-GATA3+ of Naïve CD4 T cells from Tet2+/+ and Tet2-/- cultured for 4 days in Th2-polarizing conditions (anti-CD3e, anti-CD28, anti-IFNg, recombinant IL-4 and recombinant IL-2). G, schematic describing co-culture experiments of congenically marked (Fig 3G) or 4Get (Fig 3H) Tet2+/+ naïve CD4 T cells with Tet2+/+ or Tet2-/- naïve CD4 T cells. Statistical analysis was performed by two-sided Student’s t-test in (A,B,C). p-values: * (<0.05) **(<0.01) *** (<0.001) **** (<0.0001). All panels representative of at least 2 independent experiments.

**Supplemental Figure 4, related to Figure 4:**
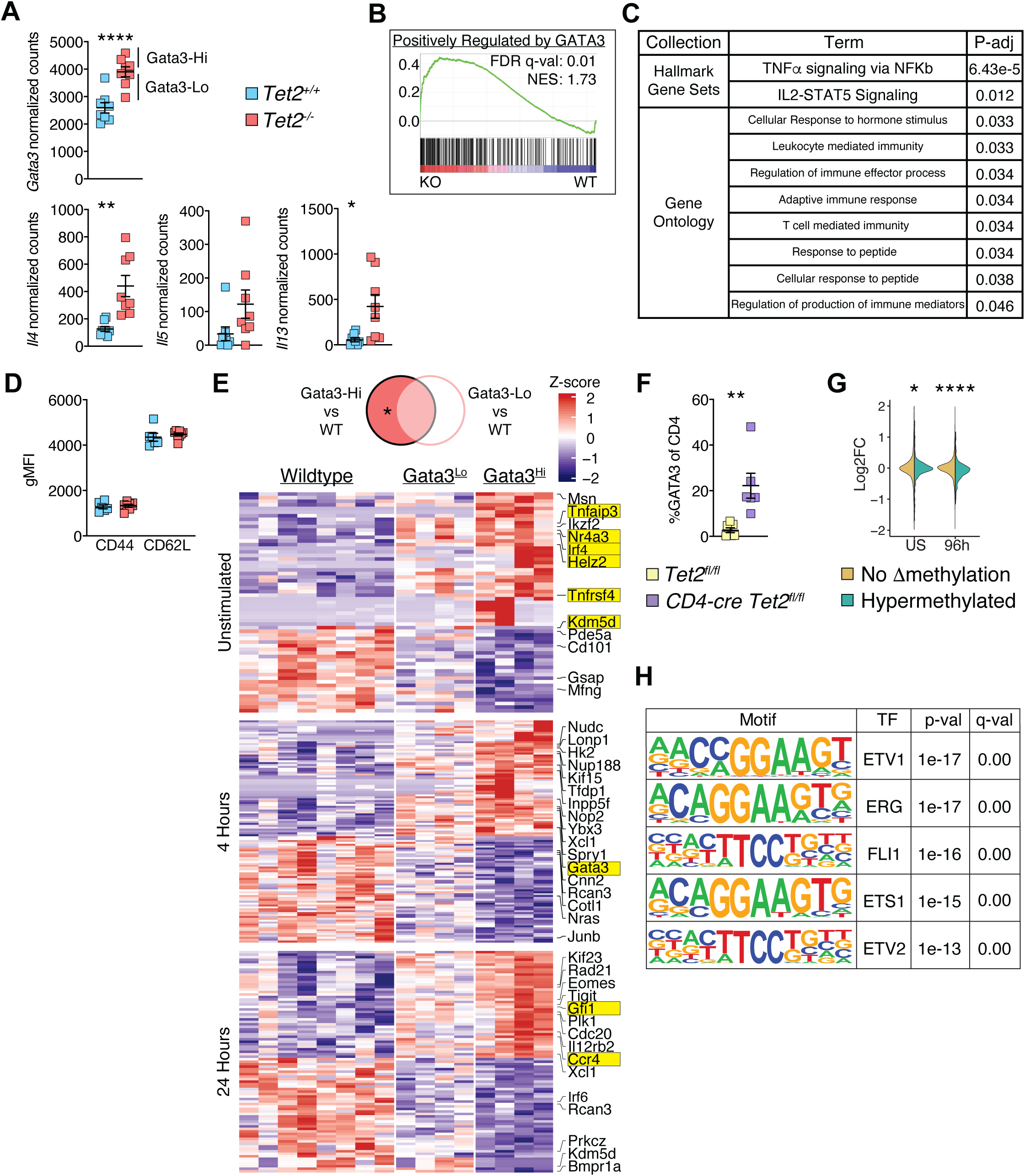
A, Normalized counts of selected signature Th2-lineage genes from RNAseq expression data of naïve CD4 T cells cultured for 4 days in non-polarizing conditions with asterisks noting adjusted p-values for each gene from the Wald test with Benjamini and Hochberg correction of p-values. B, Gene set enrichment analysis plot of naïve CD4 T cells cultured for 4 days in non-polarizing conditions. Gene set is composed of GATA3-dependent genes in Th2 cells derived in vitro^32^. C, Significant (p-adj< 0.05) gene set enrichment analysis results of sets from various collections in Tet2-deficient naïve CD4 T cells when compared to Tet2-sufficient naïve CD4 T cells. D, Expression of naïve CD4 T cell markers on freshly isolated Tet2+/+ and Tet2-/- CD4 T cells from the spleens of mice aged 6-8 weeks old. Assessed by flow cytometry and geometric mean fluorescent intensity of cells gated as naïve CD4 T cells (gated Live CD45+ CD4+ CD8a-CD62L+ CD44lo). E, Z-score of normalized expression of differentially expressed genes in Gata3Hi Tet2-deficient samples but not Gata3Lo Tet2-deficient samples (indicated by *) relative to Tet2-sufficient CD4 T cells cultured for indicated time points under non-polarizing conditions. F, Proportion of GATA3+ Foxp3- of CD4 from Naive CD4 cells from Tet2 fl/fl or CD4-cre+ Tet2 fl/fl mice cultured non-polarizing conditions for 96h. Two-sided student’s t test. G, Genes were considered if they were expressed in our Bulk RNAseq dataset and contained nearby loci for which methylation was measured in our DNA methylation array. Log 2 Fold change was plotted of genes in Tet2-deficient compared to Tet2-sufficient CD4 T cells from unstimulated naïve CD4 T cells or naïve CD4 T cells cultured under non-polarizing conditions for 96 hours. These genes were subset by whether they significantly hypermethylated loci in naïve Tet2-deficient CD4 T cells relative to Tet2-sufficient naïve CD4 T cells and a one-sided t test was used to compare the distribution of log 2 fold change values. H, Enrichment of known motifs from CHIP-seq datasets^82^ in the sequences of regions less accessible in Tet2-deficient naive CD4 T cells along with p-values representing enrichment along with q-value. p-values: * (<0.05) **(<0.01) *** (<0.001) **** (<0.0001).

**Supplemental Figure 5, related to Figure 5:**
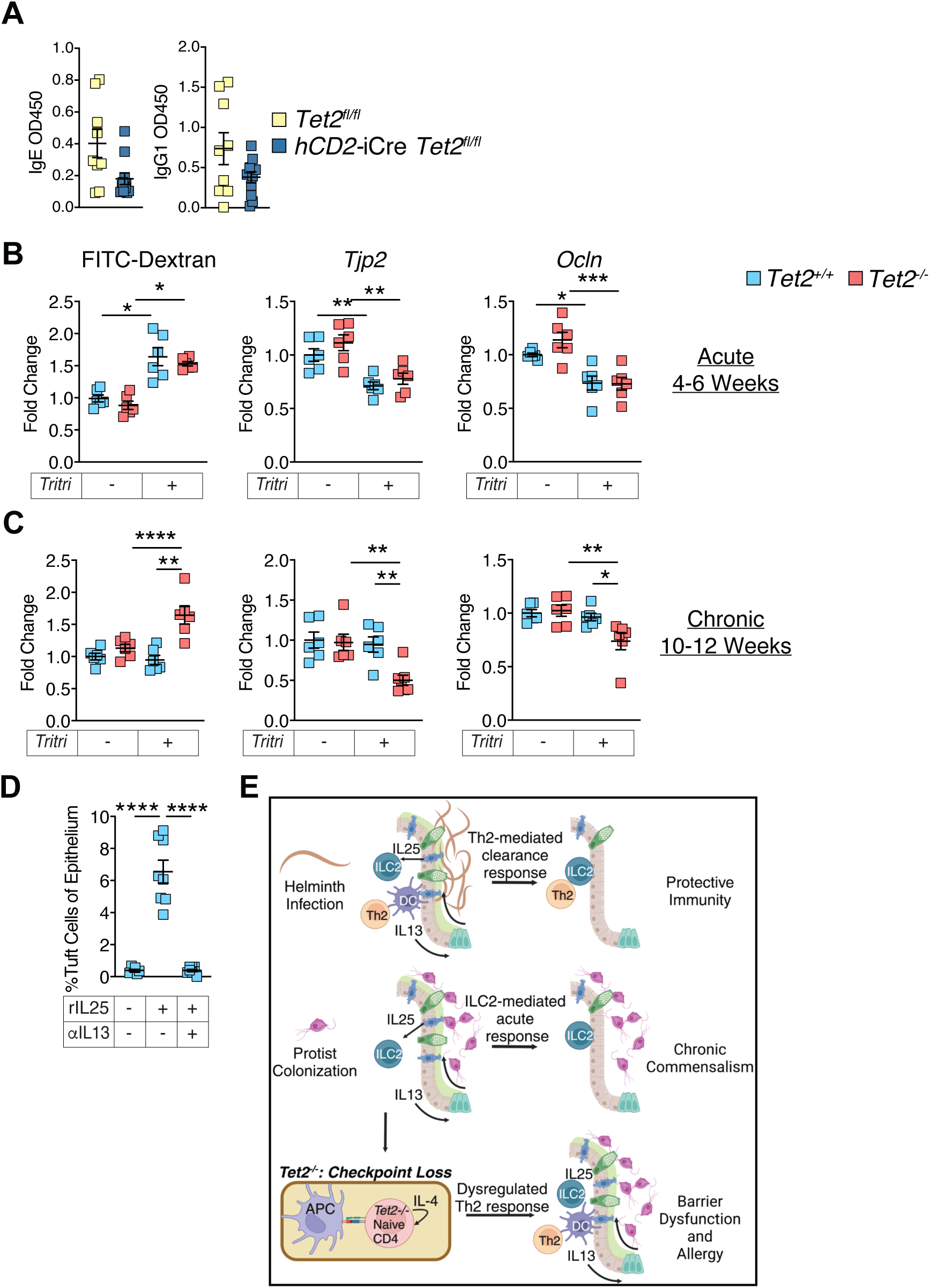
A, ELISA quantification of peanut-specific IgE and IgG1 antibodies in the serum of hCD2-Cre Tet2fl/fl mice and littermates on day 28 of sensitization prior to challenge. B/C, FITC-dextran permeability measured in the peripheral blood 3h after gavage of 0.1mg/g FITC-dextran. Represented as fold-change over littermate uncolonized Tet2+/+ mice (left). Quantitative PCR (right) of Tjp2 and Ocln measured relative to Gapdh and represented as fold-change over littermate uncolonized Tet2+/+ mice. FITC dextran and qPCR performed after acute colonization (4-6 weeks, B) and chronic colonization (10-12 weeks, C). D, Proportion of SiglecF+ CD24+ Tuft cells of Live Epcam+ epithelial cells in the epithelial fraction from the jejunum of mice treated with vehicle, recombinant IL25 or recombinant IL25 and anti-IL13 for 2 weeks. E, model representing strict control of intestinal responses to helminths and commensals and the consequences of unrestrained Th2 activation in response to chronically colonized commensal microbes, such as Tritrichomonas. Statistical analysis was performed by two-sided Student’s t-test in (A), and one-way ANOVA with Tukey multiple-comparison test in (C,D). p-values: * (<0.05) **(<0.01) *** (<0.001) **** (<0.0001)

## References

1. Tang, M.L.K., and Mullins, R.J. (2017). Food allergy: is prevalence increasing? Internal Medicine Journal 47, 256–261. 10.1111/imj.13362.

2. Anthony, R.M., Rutitzky, L.I., Urban, J.F., Jr., Stadecker, M.J., and Gause, W.C. (2007). Protective immune mechanisms in helminth infection. Nat Rev Immunol 7, 975–987. 10.1038/nri2199.

3. Blumenthal, M.N. (2012). Genetic, epigenetic, and environmental factors in asthma and allergy. Ann Allergy Asthma Immunol 108, 69–73. 10.1016/j.anai.2011.12.003.

4. Virolainen, S.J., VonHandorf, A., Viel, K., Weirauch, M.T., and Kottyan, L.C. (2023). Gene-environment interactions and their impact on human health. Genes Immun 24, 1–11. 10.1038/s41435-022-00192-6.

5. Zhu, J. (2015). T helper 2 (Th2) cell differentiation, type 2 innate lymphoid cell (ILC2) development and regulation of interleukin-4 (IL-4) and IL-13 production. Cytokine 75, 14–24. 10.1016/J.CYTO.2015.05.010.

6. Yoshimoto, T. (2018). The Hunt for the Source of Primary Interleukin-4: How We Discovered That Natural Killer T Cells and Basophils Determine T Helper Type 2 Cell Differentiation In Vivo. Front Immunol 9, 716. 10.3389/fimmu.2018.00716.

7. Ruterbusch, M., Pruner, K.B., Shehata, L., and Pepper, M. (2020). In Vivo CD4+ T Cell Differentiation and Function: Revisiting the Th1/Th2 Paradigm. 10.1146/annurev-immunol-103019-085803 38, 705–725. 10.1146/ANNUREV-IMMUNOL-103019-085803.

8. Lloyd, C.M., and Snelgrove, R.J. (2018). Type 2 immunity: Expanding our view. Sci Immunol 3. 10.1126/sciimmunol.aat1604.

9. Negrao-Correa, D., Pinho, V., Souza, D.G., Pereira, A.T., Fernandes, A., Scheuermann, K., Souza, A.L., and Teixeira, M.M. (2006). Expression of IL-4 receptor on non-bone marrow-derived cells is necessary for the timely elimination of Strongyloides venezuelensis in mice, but not for intestinal IL-4 production. Int J Parasitol 36, 1185–1195. 10.1016/j.ijpara.2006.05.005.

10. Phillips, C., Coward, W.R., Pritchard, D.I., and Hewitt, C.R. (2003). Basophils express a type 2 cytokine profile on exposure to proteases from helminths and house dust mites. J Leukoc Biol 73, 165–171. 10.1189/jlb.0702356.

11. Ivanov, I.I., Atarashi, K., Manel, N., Brodie, E.L., Shima, T., Karaoz, U., Wei, D., Goldfarb, K.C., Santee, C.A., Lynch, S.V., et al. (2009). Induction of Intestinal Th17 Cells by Segmented Filamentous Bacteria. Cell 139, 485–498. 10.1016/J.CELL.2009.09.033.

12. Nagayama, M., Yano, T., Atarashi, K., Tanoue, T., Sekiya, M., Kobayashi, Y., Sakamoto, H., Miura, K., Sunada, K., Kawaguchi, T., et al. (2020). TH1 cell-inducing Escherichia coli strain identified from the small intestinal mucosa of patients with Crohn’s disease. Gut Microbes 12, 1788898. 10.1080/19490976.2020.1788898.

13. Ivanov, I.I., Tuganbaev, T., Skelly, A.N., and Honda, K. (2022). T Cell Responses to the Microbiota. 10.1146/annurev-immunol-101320-011829 40, 559–587. 10.1146/ANNUREV-IMMUNOL-101320-011829.

14. Howitt, M.R., Lavoie, S., Michaud, M., Blum, A.M., Tran, S.V., Weinstock, J.V., Gallini, C.A., Redding, K., Margolskee, R.F., Osborne, L.C., et al. (2016). Tuft cells, taste-chemosensory cells, orchestrate parasite type 2 immunity in the gut. Science 351, 1329–1333. 10.1126/SCIENCE.AAF1648/SUPPL_FILE/AAF1648-HOWITT-SM.PDF.

15. Nadjsombati, M.S., McGinty, J.W., Lyons-Cohen, M.R., Jaffe, J.B., DiPeso, L., Schneider, C., Miller, C.N., Pollack, J.L., Nagana Gowda, G.A., Fontana, M.F., et al. (2018). Detection of Succinate by Intestinal Tuft Cells Triggers a Type 2 Innate Immune Circuit. Immunity 49, 33–41.e37. 10.1016/J.IMMUNI.2018.06.016.

16. Schneider, C., O’Leary, C.E., von Moltke, J., Liang, H.E., Ang, Q.Y., Turnbaugh, P.J., Radhakrishnan, S., Pellizzon, M., Ma, A., and Locksley, R.M. (2018). A Metabolite-Triggered Tuft Cell-ILC2 Circuit Drives Small Intestinal Remodeling. Cell 174, 271–284.e214. 10.1016/J.CELL.2018.05.014.

17. Chudnovskiy, A., Mortha, A., Kana, V., Kennard, A., Ramirez, J.D., Rahman, A., Remark, R., Mogno, I., Ng, R., Gnjatic, S., et al. (2016). Host-Protozoan Interactions Protect from Mucosal Infections through Activation of the Inflammasome. Cell 167, 444–456.e414. 10.1016/J.CELL.2016.08.076.

18. Vacca, F., and Le Gros, G. (2022). Tissue-specific immunity in helminth infections. Mucosal Immunol 15, 1212–1223. 10.1038/s41385-022-00531-w.

19. Von Moltke, J., Ji, M., Liang, H.E., and Locksley, R.M. (2015). Tuft-cell-derived IL-25 regulates an intestinal ILC2–epithelial response circuit. Nature 2015 529:7585 529, 221–225. 10.1038/nature16161.

20. Medina Sanchez, L., Siller, M., Zeng, Y., Brigleb, P.H., Sangani, K.A., Soto, A.S., Engl, C., Laughlin, C.R., Rana, M., Van Der Kraak, L., et al. (2023). The gut protist Tritrichomonas arnold restrains virus-mediated loss of oral tolerance by modulating dietary antigen-presenting dendritic cells. Immunity 56, 1862–1875.e1869. 10.1016/j.immuni.2023.06.022.

21. Jones, P.A. (2012). Functions of DNA methylation: islands, start sites, gene bodies and beyond. Nat Rev Genet 13, 484–492. 10.1038/nrg3230.

22. Li, J., Li, L., Sun, X., Deng, T., Huang, G., Li, X., Xie, Z., and Zhou, Z. (2021). Role of Tet2 in Regulating Adaptive and Innate Immunity. Frontiers in Cell and Developmental Biology 9, 1573–1573. 10.3389/FCELL.2021.665897/BIBTEX.

23. Venugopal, K., Feng, Y., Shabashvili, D., and Guryanova, O.A. (2021). Alterations to DNMT3A in Hematologic Malignancies. Cancer Res 81, 254–263. 10.1158/0008-5472.CAN-20-3033.

24. Bowman, R.L., Busque, L., and Levine, R.L. (2018). Clonal Hematopoiesis and Evolution to Hematopoietic Malignancies. Cell Stem Cell 22, 157–170. 10.1016/j.stem.2018.01.011.

25. Bowman, R.L., and Levine, R.L. (2017). TET2 in Normal and Malignant Hematopoiesis. Cold Spring Harb Perspect Med 7. 10.1101/cshperspect.a026518.

26. Meisel, M., Hinterleitner, R., Pacis, A., Chen, L., Earley, Z.M., Mayassi, T., Pierre, J.F., Ernest, J.D., Galipeau, H.J., Thuille, N., et al. (2018). Microbial signals drive pre-leukaemic myeloproliferation in a Tet2-deficient host. Nature 557, 580–584. 10.1038/s41586-018-0125-z.

27. Franzén, O., Gan, L.M., and Björkegren, J.L.M. (2019). PanglaoDB: a web server for exploration of mouse and human single-cell RNA sequencing data. Database 2019, 46–46. 10.1093/DATABASE/BAZ046.

28. Schneider, C., Lee, J., Koga, S., Ricardo-Gonzalez, R.R., Nussbaum, J.C., Smith, L.K., Villeda, S.A., Liang, H.E., and Locksley, R.M. (2019). Tissue-Resident Group 2 Innate Lymphoid Cells Differentiate by Layered Ontogeny and In Situ Perinatal Priming. Immunity 50, 1425–1438.e1425. 10.1016/J.IMMUNI.2019.04.019.

29. Roach, P.D., Wallis, P.M., and Olson, M.E. (1988). The use of metronidazole, tinidazole and dimetridazole in eliminating trichomonads from laboratory mice. Laboratory Animals 22, 361–364. 10.1258/002367788780746287.

30. Testi, R., Phillips, J.H., and Lanier, L.L. (1989). Leu 23 induction as an early marker of functional CD3/T cell antigen receptor triggering. Requirement for receptor cross-linking, prolonged elevation of intracellular [Ca++] and stimulation of protein kinase C. J Immunol 142, 1854–1860.

31. Mohrs, M., Shinkai, K., Mohrs, K., and Locksley, R.M. (2001). Analysis of type 2 immunity in vivo with a bicistronic IL-4 reporter. Immunity 15, 303–311. 10.1016/s1074-7613(01)00186-8.

32. Wei, G., Abraham, B.J., Yagi, R., Jothi, R., Cui, K., Sharma, S., Narlikar, L., Northrup, D.L., Tang, Q., Paul, W.E., et al. (2011). Genome-wide Analyses of Transcription Factor GATA3-Mediated Gene Regulation in Distinct T Cell Types. Immunity 35, 299–311. 10.1016/J.IMMUNI.2011.08.007.

33. Liberzon, A., Birger, C., Thorvaldsdottir, H., Ghandi, M., Mesirov, J.P., and Tamayo, P. (2015). The Molecular Signatures Database (MSigDB) hallmark gene set collection. Cell Syst 1, 417–425. 10.1016/j.cels.2015.12.004.

34. Ashburner, M., Ball, C.A., Blake, J.A., Botstein, D., Butler, H., Cherry, J.M., Davis, A.P., Dolinski, K., Dwight, S.S., Eppig, J.T., et al. (2000). Gene ontology: tool for the unification of biology. The Gene Ontology Consortium. Nat Genet 25, 25–29. 10.1038/75556.

35. Oh, H., and Ghosh, S. (2013). NF-kappaB: roles and regulation in different CD4(+) T-cell subsets. Immunol Rev 252, 41–51. 10.1111/imr.12033.

36. Zhu, J., Cote-Sierra, J., Guo, L., and Paul, W.E. (2003). Stat5 Activation Plays a Critical Role in Th2 Differentiation. Immunity 19, 739–748. 10.1016/S1074-7613(03)00292-9.

37. Zhu, J., Yamane, H., and Paul, W.E. (2010). Differentiation of effector CD4 T cell populations (*). Annu Rev Immunol 28, 445–489. 10.1146/annurev-immunol-030409-101212.

38. Bocek, P., Jr., Foucras, G., and Paul, W.E. (2004). Interferon gamma enhances both in vitro and in vivo priming of CD4+ T cells for IL-4 production. J Exp Med 199, 1619–1630. 10.1084/jem.20032014.

39. Torres, K.C., Dutra, W.O., and Gollob, K.J. (2004). Endogenous IL-4 and IFN-gamma are essential for expression of Th2, but not Th1 cytokine message during the early differentiation of human CD4+ T helper cells. Hum Immunol 65, 1328–1335. 10.1016/j.humimm.2004.06.007.

40. Porter, C.M., and Clipstone, N.A. (2002). Sustained NFAT signaling promotes a Th1-like pattern of gene expression in primary murine CD4+ T cells. J Immunol 168, 4936–4945. 10.4049/jimmunol.168.10.4936.

41. Rengarajan, J., Tang, B., and Glimcher, L.H. (2002). NFATc2 and NFATc3 regulate T(H)2 differentiation and modulate TCR-responsiveness of naive T(H)cells. Nat Immunol 3, 48–54. 10.1038/ni744.

42. Fang, D., Cui, K., Hu, G., Gurram, R.K., Zhong, C., Oler, A.J., Yagi, R., Zhao, M., Sharma, S., Liu, P., et al. (2018). Bcl11b, a novel GATA3-interacting protein, suppresses Th1 while limiting Th2 cell differentiation. J Exp Med 215, 1449–1462. 10.1084/jem.20171127.

43. Komine, O., Hayashi, K., Natsume, W., Watanabe, T., Seki, Y., Seki, N., Yagi, R., Sukzuki, W., Tamauchi, H., Hozumi, K., et al. (2003). The Runx1 transcription factor inhibits the differentiation of naive CD4+ T cells into the Th2 lineage by repressing GATA3 expression. J Exp Med 198, 51–61. 10.1084/jem.20021200.

44. Bachus, H., McLaughlin, E., Lewis, C., Papillion, A.M., Benveniste, E.N., Hill, D.D., Rosenberg, A.F., Ballesteros-Tato, A., and Leon, B. (2023). IL-6 prevents Th2 cell polarization by promoting SOCS3-dependent suppression of IL-2 signaling. Cell Mol Immunol 20, 651–665. 10.1038/s41423-023-01012-1.

45. Tangye, S.G., Pillay, B., Randall, K.L., Avery, D.T., Phan, T.G., Gray, P., Ziegler, J.B., Smart, J.M., Peake, J., Arkwright, P.D., et al. (2017). Dedicator of cytokinesis 8-deficient CD4(+) T cells are biased to a T(H)2 effector fate at the expense of T(H)1 and T(H)17 cells. J Allergy Clin Immunol 139, 933–949. 10.1016/j.jaci.2016.07.016.

46. Hossain, M.B., Hosokawa, H., Hasegawa, A., Watarai, H., Taniguchi, M., Yamashita, M., and Nakayama, T. (2008). Lymphoid enhancer factor interacts with GATA-3 and controls its function in T helper type 2 cells. Immunology 125, 377–386. 10.1111/j.1365-2567.2008.02854.x.

47. Kuwahara, M., Ise, W., Ochi, M., Suzuki, J., Kometani, K., Maruyama, S., Izumoto, M., Matsumoto, A., Takemori, N., Takemori, A., et al. (2016). Bach2-Batf interactions control Th2-type immune response by regulating the IL-4 amplification loop. Nat Commun 7, 12596. 10.1038/ncomms12596.

48. Kitamura, F., Kitamura, N., Mori, A., Tatsumi, H., Nemoto, S., Miyoshi, H., Miyatake, S., Hiroi, T., and Kaminuma, O. (2010). Selective down-regulation of Th2 cytokines by C-terminal binding protein 2 in human T cells. Int Arch Allergy Immunol 152 *Suppl 1*, 18–21. 10.1159/000312121.

49. Kim, C.J., Lee, C.G., Jung, J.Y., Ghosh, A., Hasan, S.N., Hwang, S.M., Kang, H., Lee, C., Kim, G.C., Rudra, D., et al. (2018). The Transcription Factor Ets1 Suppresses T Follicular Helper Type 2 Cell Differentiation to Halt the Onset of Systemic Lupus Erythematosus. Immunity 49, 1034–1048 e1038. 10.1016/j.immuni.2018.10.012.

50. Hebenstreit, D., Giaisi, M., Treiber, M.K., Zhang, X.B., Mi, H.F., Horejs-Hoeck, J., Andersen, K.G., Krammer, P.H., Duschl, A., and Li-Weber, M. (2008). LEF-1 negatively controls interleukin-4 expression through a proximal promoter regulatory element. J Biol Chem 283, 22490–22497. 10.1074/jbc.M804096200.

51. San Luis, B., Sondgeroth, B., Nassar, N., and Carpino, N. (2011). Sts-2 is a phosphatase that negatively regulates zeta-associated protein (ZAP)-70 and T cell receptor signaling pathways. J Biol Chem 286, 15943–15954. 10.1074/jbc.M110.177634.

52. Sofi, M.H., Heinrichs, J., Dany, M., Nguyen, H., Dai, M., Bastian, D., Schutt, S., Wu, Y., Daenthanasanmak, A., Gencer, S., et al. (2017). Ceramide synthesis regulates T cell activity and GVHD development. JCI Insight 2. 10.1172/jci.insight.91701.

53. Collier, C., Wucherer, K., McWhorter, M., Jenkins, C., Bartlett, A., Roychoudhuri, R., and Eil, R. (2024). Intracellular K+ Limits T-cell Exhaustion and Preserves Antitumor Function. Cancer Immunol Res 12, 36–47. 10.1158/2326-6066.CIR-23-0319.

54. Yuan, Y., Yang, L., Liu, T., Zhang, H., and Lu, Q. (2019). Osteoclastogenesis inhibition by mutated IGSF23 results in human osteopetrosis. Cell Prolif 52, e12693. 10.1111/cpr.12693.

55. Richert-Spuhler, L.E., Mar, C.M., Shinde, P., Wu, F., Hong, T., Greene, E., Hou, S., Thomas, K., Gottardo, R., Mugo, N., et al. (2021). CD101 genetic variants modify regulatory and conventional T cell phenotypes and functions. Cell Rep Med 2, 100322. 10.1016/j.xcrm.2021.100322.

56. Schey, R., Dornhoff, H., Baier, J.L., Purtak, M., Opoka, R., Koller, A.K., Atreya, R., Rau, T.T., Daniel, C., Amann, K., et al. (2016). CD101 inhibits the expansion of colitogenic T cells. Mucosal Immunol 9, 1205–1217. 10.1038/mi.2015.139.

57. Shifrut, E., Carnevale, J., Tobin, V., Roth, T.L., Woo, J.M., Bui, C.T., Li, P.J., Diolaiti, M.E., Ashworth, A., and Marson, A. (2018). Genome-wide CRISPR Screens in Primary Human T Cells Reveal Key Regulators of Immune Function. Cell 175, 1958–1971 e1915. 10.1016/j.cell.2018.10.024.

58. Ta, H.M., Roy, D., Zhang, K., Alban, T., Juric, I., Dong, J., Parthasarathy, P.B., Patnaik, S., Delaney, E., Gilmour, C., et al. (2024). LRIG1 engages ligand VISTA and impairs tumor-specific CD8(+) T cell responses. Sci Immunol 9, eadi7418. 10.1126/sciimmunol.adi7418.

59. Baessler, A., Novis, C.L., Shen, Z., Perovanovic, J., Wadsworth, M., Thiede, K.A., Sircy, L.M., Harrison-Chau, M., Nguyen, N.X., Varley, K.E., et al. (2022). Tet2 coordinates with Foxo1 and Runx1 to balance T follicular helper cell and T helper 1 cell differentiation. Sci Adv 8, eabm4982. 10.1126/sciadv.abm4982.

60. Endo, Y., Hirahara, K., Iinuma, T., Shinoda, K., Tumes, D.J., Asou, H.K., Matsugae, N., Obata-Ninomiya, K., Yamamoto, H., Motohashi, S., et al. (2015). The interleukin-33-p38 kinase axis confers memory T helper 2 cell pathogenicity in the airway. Immunity 42, 294–308. 10.1016/j.immuni.2015.01.016.

61. Lee, J.B., Chen, C.Y., Liu, B., Mugge, L., Angkasekwinai, P., Facchinetti, V., Dong, C., Liu, Y.J., Rothenberg, M.E., Hogan, S.P., et al. (2016). IL-25 and CD4+ TH2 cells enhance type 2 innate lymphoid cell–derived IL-13 production, which promotes IgE-mediated experimental food allergy. Journal of Allergy and Clinical Immunology 137, 1216–1225.e1215. 10.1016/J.JACI.2015.09.019.

62. Smit, J.J., Willemsen, K., Hassing, I., Fiechter, D., Storm, G., van Bloois, L., Leusen, J.H., Pennings, M., Zaiss, D., and Pieters, R.H. (2011). Contribution of classic and alternative effector pathways in peanut-induced anaphylactic responses. PLoS One 6, e28917. 10.1371/journal.pone.0028917.

63. Samadi, N., Klems, M., and Untersmayr, E. (2018). The role of gastrointestinal permeability in food allergy. Annals of Allergy, Asthma & Immunology 121, 168–173. 10.1016/J.ANAI.2018.05.010.

64. Horowitz, A., Chanez-Paredes, S.D., Haest, X., and Turner, J.R. (2023). Paracellular permeability and tight junction regulation in gut health and disease. Nat Rev Gastroenterol Hepatol 20, 417–432. 10.1038/s41575-023-00766-3.

65. Madden, K.B., Yeung, K.A., Zhao, A., Gause, W.C., Finkelman, F.D., Katona, I.M., Urban, J.F., Jr., and Shea-Donohue, T. (2004). Enteric nematodes induce stereotypic STAT6-dependent alterations in intestinal epithelial cell function. J Immunol 172, 5616–5621. 10.4049/jimmunol.172.9.5616.

66. Shea-Donohue, T., Sullivan, C., Finkelman, F.D., Madden, K.B., Morris, S.C., Goldhill, J., Pineiro-Carrero, V., and Urban, J.F., Jr. (2001). The role of IL-4 in Heligmosomoides polygyrus-induced alterations in murine intestinal epithelial cell function. J Immunol 167, 2234–2239. 10.4049/jimmunol.167.4.2234.

67. Moyat, M., Lebon, L., Perdijk, O., Wickramasinghe, L.C., Zaiss, M.M., Mosconi, I., Volpe, B., Guenat, N., Shah, K., Coakley, G., et al. (2022). Microbial regulation of intestinal motility provides resistance against helminth infection. Mucosal Immunol 15, 1283–1295. 10.1038/s41385-022-00498-8.

68. Lin, Y., Li, B., Yang, X., Liu, T., Shi, T., Deng, B., Zhang, Y., Jia, L., Jiang, Z., and He, R. (2019). Non-hematopoietic STAT6 induces epithelial tight junction dysfunction and promotes intestinal inflammation and tumorigenesis. Mucosal Immunol 12, 1304–1315. 10.1038/s41385-019-0204-y.

69. Gerbe, F., Sidot, E., Smyth, D.J., Ohmoto, M., Matsumoto, I., Dardalhon, V., Cesses, P., Garnier, L., Pouzolles, M., Brulin, B., et al. (2016). Intestinal epithelial tuft cells initiate type 2 mucosal immunity to helminth parasites. Nature 2016 529:7585 529, 226–230. 10.1038/nature16527.

70. van Panhuys, N., Prout, M., Forbes, E., Min, B., Paul, W.E., and Le Gros, G. (2011). Basophils are the major producers of IL-4 during primary helminth infection. J Immunol 186, 2719–2728. 10.4049/jimmunol.1000940.

71. Min, B., Prout, M., Hu-Li, J., Zhu, J., Jankovic, D., Morgan, E.S., Urban, J.F., Dvorak, A.M., Finkelman, F.D., LeGros, G., and Paul, W.E. (2004). Basophils Produce IL-4 and Accumulate in Tissues after Infection with a Th2-inducing Parasite. Journal of Experimental Medicine 200, 507–517. 10.1084/JEM.20040590.

72. Johansson, E., and Mersha, T.B. (2021). Genetics of Food Allergy. Immunol Allergy Clin North Am 41, 301–319. 10.1016/j.iac.2021.01.010.

73. Yagi, J., Arimura, Y., Takatori, H., Nakajima, H., Iwamoto, I., and Uchiyama, T. (2006). Genetic background influences Th cell differentiation by controlling the capacity for IL-2-induced IL-4 production by naive CD4+ T cells. International Immunology 18, 1681–1690. 10.1093/INTIMM/DXL102.

74. Yagi, R., Suzuki, W., Seki, N., Kohyama, M., Inoue, T., Arai, T., and Kubo, M. (2002). The IL-4 production capability of different strains of naive CD4(+) T cells controls the direction of the T(h) cell response. Int Immunol 14, 1–11. 10.1093/intimm/14.1.1.

75. Howitt, M.R., Cao, Y.G., Gologorsky, M.B., Li, J.A., Haber, A.L., Biton, M., Lang, J., Michaud, M., Regev, A., and Garrett, W.S. (2020). The Taste Receptor TAS1R3 Regulates Small Intestinal Tuft Cell Homeostasis. ImmunoHorizons 4, 23–32. 10.4049/IMMUNOHORIZONS.1900099/-/DCSUPPLEMENTAL.

76. Burke, M.J., and Zalewska-Szewczyk, B. (2022). Hypersensitivity reactions to asparaginase therapy in acute lymphoblastic leukemia: immunology and clinical consequences. Future Oncol 18, 1285–1299. 10.2217/fon-2021-1288.

77. Tan, L., Fu, L., Zheng, L., Fan, W., Tan, H., Tao, Z., and Xu, Y. (2022). TET2 Regulates 5-Hydroxymethylcytosine Signature and CD4(+) T-Cell Balance in Allergic Rhinitis. Allergy Asthma Immunol Res 14, 254–272. 10.4168/aair.2022.14.2.254.

78. Cardenas, A., Fadadu, R.P., and Koppelman, G.H. (2023). Epigenome-wide association studies of allergic disease and the environment. J Allergy Clin Immunol 152, 582–590. 10.1016/j.jaci.2023.05.020.

79. Peng, V., Xing, X., Bando, J.K., Trsan, T., Di Luccia, B., Collins, P.L., Li, D., Wang, W.L., Lee, H.J., Oltz, E.M., et al. (2022). Whole-genome profiling of DNA methylation and hydroxymethylation identifies distinct regulatory programs among innate lymphocytes. Nature Immunology 2022 23:4 23, 619–631. 10.1038/s41590-022-01164-8.

80. van Breugel, M., Qi, C., Xu, Z., Pedersen, C.T., Petoukhov, I., Vonk, J.M., Gehring, U., Berg, M., Bugel, M., Carpaij, O.A., et al. (2022). Nasal DNA methylation at three CpG sites predicts childhood allergic disease. Nat Commun 13, 7415. 10.1038/s41467-022-35088-6.

81. Xu, C.J., Gruzieva, O., Qi, C., Esplugues, A., Gehring, U., Bergstrom, A., Mason, D., Chatzi, L., Porta, D., Lodrup Carlsen, K.C., et al. (2021). Shared DNA methylation signatures in childhood allergy: The MeDALL study. J Allergy Clin Immunol 147, 1031–1040. 10.1016/j.jaci.2020.11.044.

82. Heinz, S., Benner, C., Spann, N., Bertolino, E., Lin, Y.C., Laslo, P., Cheng, J.X., Murre, C., Singh, H., and Glass, C.K. (2010). Simple combinations of lineage-determining transcription factors prime cis-regulatory elements required for macrophage and B cell identities. Mol Cell 38, 576–589. 10.1016/j.molcel.2010.05.004.

83. Ferrer-Font, L., Mehta, P., Harmos, P., Schmidt, A.J., Chappell, S., Price, K.M., Hermans, I.F., Ronchese, F., le Gros, G., and Mayer, J.U. (2020). High-dimensional analysis of intestinal immune cells during helminth infection. Elife 9. 10.7554/eLife.51678.

84. Esterházy, D., Canesso, M.C.C., Mesin, L., Muller, P.A., de Castro, T.B.R., Lockhart, A., ElJalby, M., Faria, A.M.C., and Mucida, D. (2019). Compartmentalized gut lymph node drainage dictates adaptive immune responses. Nature 2019 569:7754 569, 126–130. 10.1038/s41586-019-1125-3.

85. Nik, A.M., and Carlsson, P. (2013). Separation of intact intestinal epithelium from mesenchyme. Biotechniques 55, 42–44. 10.2144/000114055.

86. Grandi, F.C., Modi, H., Kampman, L., and Corces, M.R. (2022). Chromatin accessibility profiling by ATAC-seq. Nat Protoc 17, 1518–1552. 10.1038/s41596-022-00692-9.

87. Martino, D., Neeland, M., Dang, T., Cobb, J., Ellis, J., Barnett, A., Tang, M., Vuillermin, P., Allen, K., and Saffery, R. (2018). Epigenetic dysregulation of naive CD4+ T-cell activation genes in childhood food allergy. Nat Commun 9, 3308. 10.1038/s41467-01805608-4.

88. Subramanian, A., Tamayo, P., Mootha, V.K., Mukherjee, S., Ebert, B.L., Gillette, M.A., Paulovich, A., Pomeroy, S.L., Golub, T.R., Lander, E.S., and Mesirov, J.P. (2005). Gene set enrichment analysis: A knowledge-based approach for interpreting genome-wide expression profiles. Proceedings of the National Academy of Sciences of the United States of America 102, 15545–15550. 10.1073/PNAS.0506580102/SUPPL_FILE/06580FIG7.JPG.

89. Patel, H.E.-C., Jose; Langer, Björn; Ewels, Phil; nf-core bot; Garcia, Maxime U; Syme, Robert; Peltzer, Alexander; Talbot, Adam; Behrens, Drew; Gabernet, Gisela; Jin, Mingda; Hörtenhubrer, Matthias; Gonzalez Rodriguez, Jonatan; Menden, Kevin; Ömer, An. (2023). nf-core/atacseq (2.1.2). 10.5281/zenodo.8222875.

90. Ritchie, M.E., Phipson, B., Wu, D., Hu, Y., Law, C.W., Shi, W., and Smyth, G.K. (2015). limma powers differential expression analyses for RNA-sequencing and microarray studies. Nucleic Acids Res 43, e47. 10.1093/nar/gkv007.

91. Ding, W., Kaur, D., Horvath, S., and Zhou, W. (2023). Comparative epigenome analysis using Infinium DNA methylation BeadChips. Brief Bioinform 24 10.1093/bib/bbac617.

92. Du, P., Zhang, X., Huang, C.C., Jafari, N., Kibbe, W.A., Hou, L., and Lin, S.M. (2010). Comparison of Beta-value and M-value methods for quantifying methylation levels by microarray analysis. BMC Bioinformatics 11, 587. 10.1186/1471-2105-11-587.

